# A Human Neuronal Cell Model of Endogenous TDP-43 A315T Reveals Altered Protein Dynamics and Disease-Relevant Cellular Dysfunction

**DOI:** 10.64898/2026.09.02.745749

**Authors:** Maria Elena Cicardi, Sara Antonini, Ivette Martorell Serra, Megan Muench, Nikolas Kinney, Amandeep Girdhar, Riccardo Cristofani, Angelo Poletti, Jared Sterneckert, Valeria Crippa, Matteo Bordi, Piera Pasinelli, Marco Tigano, Lin Guo, Davide Trotti

## Abstract

TAR DNA-binding protein 43 (TDP-43) aggregation is the defining pathological hallmark of nearly all cases of amyotrophic lateral sclerosis (ALS), yet physiologically relevant human models that faithfully recapitulate disease-associated TDP-43 proteinopathy and dysfunction remain limited. To cover this gap, we generated a novel human-based model of cortical neurons carrying the endogenous ALS-linked TDP-43 A315T mutation together with an in-frame Dendra2 fluorescent reporter, enabling temporal and spatial monitoring of the protein. Neurons expressing TDP-43 A315T exhibited progressive neurite degeneration, altered neuronal activity, and impaired mitochondrial respiration, recapitulating several ALS-associated phenotypes. Our model also displays autophagy-dependent accumulation of cytoplasmic aggregates of mutant TDP-43 without overt loss of nuclear function, maintaining normal processing of canonical cryptic exon targets. In contrast, experimental induction of TDP-43 nuclear exclusion readily triggered cryptic exon incorporation, demonstrating that the model faithfully reproduces loss-of-function phenotypes under stress conditions. In line with perturbed protein solubility, mutant TDP-43 neurons show increased stress granule (SG) formation at baseline and under an oxidative stress condition. Finally, treatment with the RNA chaperone Clip34 significantly reduced TDP-43 aggregation under both basal and oxidative stress conditions as well as its localization to SGs. Taken together, these findings establish a physiologically relevant human model that separates early TDP-43 toxic gain-of-function from basal loss-of-function while providing a robust platform for investigating TDP-43 biology and accelerating therapeutic discovery in ALS.

## Introduction

Amyotrophic lateral sclerosis (ALS) is a rapidly progressive neurodegenerative condition with no effective disease-modifying therapy. Although ALS is clinically and genetically heterogeneous, encompassing both familial and sporadic forms, the accumulation of TDP-43 inclusions in affected regions of the CNS is the defining pathological feature of nearly all cases^1^. TDP-43 is a ubiquitously expressed nuclear protein whose mislocalization to the cytoplasm, phosphorylation, truncation, and aggregation characterize postmortem ALS tissues^1,2^. TDP-43 aggregates are also a hallmark of approximately 30% of frontotemporal dementia cases^1^. TDP-43 aggregates can be detected in Alzheimer’s disease patients^3^, or develop as a consequence of conditions like traumatic brain injury (TBI)^4^; they may also be observed in the CNS during aging^5^, representing a converging pathogenic mechanism for multiple neurological chronic and acute conditions.

TDP-43 regulates the expression of numerous genes, in a spatial restricted manner. Depending on its localization TDP-43 affects different transcripts which are critical for different biological processes. First, its nuclear depletion results in aberrant splicing events, such as retention of intronic sequences that are normally not included in mature mRNA^6^. These retained sequences, known as cryptic exons (CEs), have become increasingly associated with ALS pathology^7–9^. CE insertion reduces levels of physiological transcripts, as the resulting aberrant mRNAs are rapidly eliminated through nonsense-mediated decay (NMD) and other degradation mechanisms^10,11^. Downregulation of STMN^9^ and UNC13^8^ has been linked to TDP-43 pathology across ALS and related dementias, supporting their relevance as downstream targets of TDP-43 nuclear depletion and linking TDP-43 loss of function (LOF) to neurodegeneration. CE insertion has also been observed in additional transcripts, including POLDIP3, CAMK2B, and HDGFL2, in post-mortem patient tissues and across multiple disease models^12^. Outside of the nucleus, TDP-43 associates with RNAs in specialized neuronal regions including the neurites and the mitochondria. In the neurites TDP-43 binds a large number of mRNA mediating their degradation maintaining the correct balance of gene expression^13^. In mitochondria, TDP-43 affects the stability of specific mitochondria tRNA and mRNA and miRNA, but its specific role is still matter of debate. It has been reported that TDP-43 association with specific mitochondrial mRNAs destabilizes them, disrupting the electron transport chain and impairing mitochondrial dynamics and functions^14–17^. Interestingly, it has been shown that TDP-43 binds mtRNA stabilizing it, and its depletion leads to mitochondrial dysfunction^18^. TDP-43 aggregates also directly causes mitochondrial imbalances by perturbing nuclear encoded mitochondrial gene products^19^. Notably, aberrant mitochondrial accumulation of mutant TDP-43 causes the formation of pores on the mitochondrial membrane which lead to mitochondrial DNA (mtDNA) leakage in the cytosol and innate immunity activation through cGAS-STING pathway^20^. Together, this evidence suggest that TDP-43 localization is strongly connected to its toxicity.

Mutations in TDP-43 cause familial ALS, and point mutations such as A315T alter the protein’s intrinsic propensity to misfold and form detergent- and protease-resistant inclusions^21^. Located in the C-terminal low-complexity region, the A315T mutation accelerates protein aggregation *in vitro* and *in vivo*, enhances formation of aberrant C-terminal fragments, and correlates with increased neurotoxicity^15,22–25^. *In vivo*, mice expressing human TDP-43 A315T display motor dysfunction, motor neuron loss, and pathological TDP-43 aggregation and truncation in affected regions.

TDP-43 turnover highly depends on functional degradative pathways. While the proteasome is able to degrade monomeric TDP-43 both in the cytoplasm and in the nucleus, autophagy being a uniquely cytoplasmic process, takes care of TDP-43 aggregates^26,27^; impaired autophagy indeed promotes TDP-43 accumulation^28^, whereas enhanced autophagy reduces aggregation and improves disease outcomes^29^.

Importantly, there is currently a lack of robust human models to model TDP-43 aggregation and related pathogenic events. Here, we use human-derived cortical neurons to model TDP-43-related pathological mechanisms. We reason that harnessing the known, toxic A315T mutation could recapitulate ALS-related mechanisms, which could be extended to other ALS phenotypes. Here, we built a human-based model in which we characterized several disease-associated phenotypes, such as neurite shortening, impaired cellular respiration, and changes in the cell transcriptome. We also identify a reproducible TDP-43 aggregation phenotype, which is attenuated by the novel Clip34 molecule, providing proof of concept that this model is a platform to evaluate candidate ALS therapeutics.

## Results

### Expression of TDP-43 A315T mutant impairs cortical neuron function and survival

To investigate the consequences of the ALS-associated TDP-43 A315T mutation in a physiologically relevant human model, we sought to create a cell model that more closely resembles the pathological mechanisms associated with mutant TDP-43. To do so, we introduced the mutation into a control human induced pluripotent stem cell (iPSC) line by CRISPR-Cas9-mediated genome editing. To enable longitudinal monitoring of endogenous TDP-43 dynamics, we simultaneously inserted a Dendra2 photo-switchable fluorescent reporter in-frame with C-terminal TDP-43, generating isogenic control and TDP-43 A315T iPSC lines (**Fig. S1a**).

Individual clones were initially selected based on Dendra2 fluorescence to identify successful knock-in events. Clones with comparable Dendra2 expression between the TDP-43 WT and TDP-43 A315T lines were subsequently validated by Sanger sequencing to confirm correct genome editing (**Fig. S1b**) and by G-banded karyotyping to verify genomic integrity (**Fig. S1c**). Alkaline phosphatase assay confirmed retention of pluripotency by the selected clones (**Fig. S1d**) as well as positivity to stem markers such as NANOG (**Fig. S1e**), OCT4, and SOX2 (**Fig. S1f**). Both WT and A315T lines express TDP-43-Dendra2, which is correctly localized to the nucleus (**Fig. S1f**). Next, we used the established hNGN2-BFP-Puromycin Piggyback system to induce differentiation of cortical neurons upon doxycycline addition (**Fig. S2a**). Cells that integrated the Piggyback cargo were first selected by puromycin, followed by the sorting of double-positive Dendra2/BFP cells by flow-cytometry (**Fig. S2b**). This strategy enabled enrichment of double-positive populations at ∼90% in both cell lines (**Fig. S2c**). After passaging both lines a few times to ensure loss of residual episomal plasmids, we used specific primers spanning the tetracycline-responsive element and the hNGN2 gene to verify stable insertion of the cassette into the genome (**Fig. S2d**). We assessed pluripotency retention using alkaline phosphatase assays (**Fig. S2e**) and SOX2 and OCT4 expression by IF (**Fig. S2f**). Using qPCR, we confirmed that the newly derived cell lines expressed SOX2 (**Fig. S2g**), OCT4 (**Fig S2h**), and NANOG (**Fig. S2i**) at comparable levels, while completely lacking NEUN expression (**Fig. S2j**).

To investigate the effects of the A315T mutation on the health and functionality of mature cortical neurons we established an optimized differentiation regimen of TDP-43 wild-type and A315T I3-iPSC lines by introducing a 2-day puromycin after 5 days of hNGN2 induction with doxycycline. This treatment effectively removed all NGN2-negative cells (**Fig S2k**), allowing for high-efficiency maturation of cortical neurons in 21 days *in vitro* (DIV), evidenced by positive SMI312 expression (**Fig. 1a**). Neuronal cultures are maintained in Brainphys medium, which includes glucose at 2.5 mM, closely resembling the brain’s physiological concentration.

**Figure 1.**
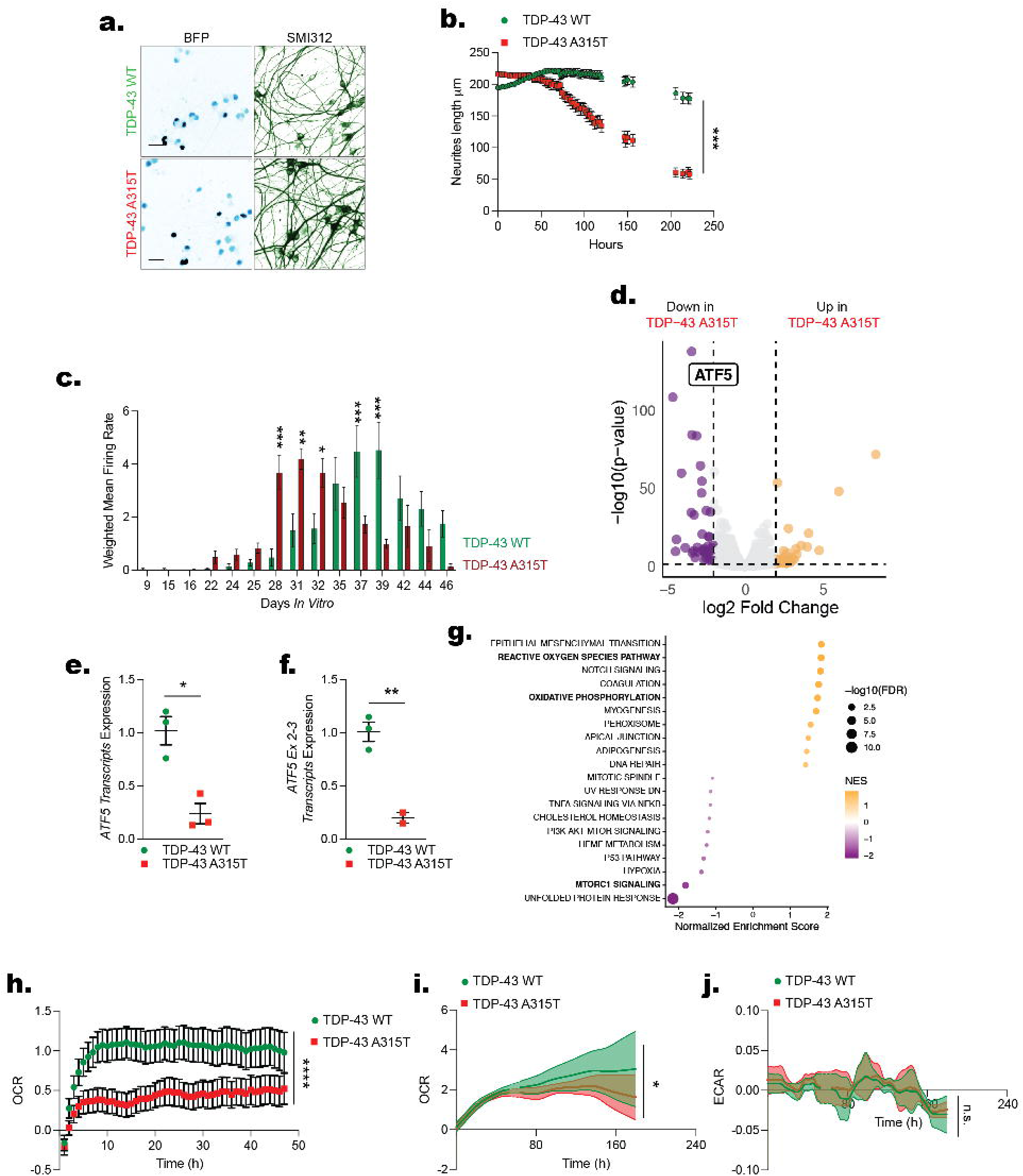
TDP-43 A315T cortical neurons display reduced neurite length, altered neuronal activity, transcriptional dysregulation, and impaired mitochondrial function. (a) Representative immunofluorescence images of mature I3 neurons from TDP-43 WT (top) and TDP-43 A315T (bottom) lines stained for SMI312 (green/dark). BFP is shown in cyan. Scale Bar = 20µm. (b) Neurite length over time (hours in vitro) in TDP-43 WT (green) and TDP-43 A315T (red) I3 neurons, quantified by live imaging. TDP-43 A315T neurons exhibit significantly shorter neurites starting from early time points. Data presented as mean ± SEM. *** p<0.001, Two-Way ANOVA. n = 8 independent replicates. (c) Weighted mean firing rate (spikes/second) measured by multielectrode array (MEA) recording over the course of neuronal maturation (Days in Vitro 9–46) in TDP-43 WT (green) and TDP-43 A315T (red) neurons. Data presented as mean ± SEM per time point. * p<0.05, ** p<0.01, *** p<0.001, Two-Way ANOVA, comparing genotypes. n = 12 independent replicates. (d) Volcano plot of differentially expressed genes (DEGs) from bulk RNA sequencing comparing TDP-43 A315T to TDP-43 WT I3 neurons. Significantly downregulated genes (left, purple) and upregulated genes (right, orange/yellow) are highlighted. Dashed lines indicate log2FC = ±1 and −log10(p-value) thresholds. (e) qRT-PCR validation of **ATF5** total transcript expression in TDP-43 WT and TDP-43 A315T neurons. Data presented as mean ± SEM, n = 2-3 independent replicates. * p<0.05, Student T-test. (f) qRT-PCR quantification of **ATF5** junction between exon 2–3 in TDP-43 WT and TDP-43 A315T neurons, further confirming downregulation at the transcript level. Data presented as mean ± SEM. ** p<0.01, Student T-test. (g) Gene Set Enrichment Analysis (GSEA) bubble plot from bulk RNAseq data. Pathways are ranked by Normalized Enrichment Score (NES). Bubble size represents −log10(FDR) and color indicates NES magnitude (scale inset). (h) OCR measurement through **Resipher** over ∼50 hours monitoring TDP-43 WT (green) and TDP-43 A315T (red) I3 neurons. **** p<0.0001, Two-Way ANOVA. n = 8 independent replicates. (i-j) **DolphinQ**-derived OCR (i) and ECAR (j) traces over ∼240 hours monitoring TDP-43 WT (green) and TDP-43 A315T (red) I3 neurons. Shaded areas represent SEM. * p<0.05, Two-Way ANOVA. n = 12 independent replicates.

Continuous brightfield live imaging showed a marked reduction in neurite length over time in mature neurons expressing TDP-43 A315T (**Fig 1b**). Loss in neurites was accompanied by aberrant neuronal activity, which we measured through multi-electrode array (MEA) recording over the course of 40 days. The two lines showed different mean firing-rate peaks, with earlier emergence of consistent firing activity in the mutant line (peak activity at DIV 28–32), whereas the control line peaked at DIV 37 and 39 (**Fig 1c**). In the mutant line, the mean firing rate decreased over time after DIV 28–32, possibly indicating neurodegeneration, while this phenomenon occurred at later time points in the wild-type line (**Fig 1c**). To assess which molecular pathways were most impacted by the presence of the mutation, we performed unbiased RNA sequencing of DIV21 neurons and analyzed differentially expressed genes (DEGs). Strikingly, we found **95** upregulated and **101** downregulated DEGs between the mutant and the wild type line. Intriguingly, we found that the top hit among the downregulated genes in the mutant line was ATF5, encoding a stress-responsive transcriptional factor activated by mitochondrial stress (**Fig 1d**). We confirmed ATF5 reduction by qPCR using two different primer sets spanning different exon junctions and observed a ∼75% reduction in ATF5 transcripts in the mutant compared to wild-type lines (**Fig 1e, f**). In support of possible mitochondrial stress downstream of TDP-43 mutation, Gene Set Enrichment Analysis (GSEA) revealed upregulation of pathways such as reactive oxygen species and oxidative phosphorylation (**Fig 1g**). Because GSEA indicated dysregulation of mitochondrial pathways, we used the Resipher technology (Lucid Scientific) to monitor the live-cell oxygen consumption rate (OCR) of wild-type and mutant TDP-43 lines over 48 hrs, starting at DIV10 (**Fig 1h**). TDP-43 A315T neurons displayed a significant decrease in OCR compared to isogenic controls, confirming mitochondrial dysfunction (**Fig 1h**). The decrease in OCR was reproduced over a longer time window with a different live-cell technology, the DolphinQ (Lead Biosystems) (**Fig 1i**). In addition to OCR, DolphinQ enables continuous recording of media pH, which can be converted to Extracellular Acidification Rate (ECAR), a standard measure of glycolytic activity. In contrast to OCR, ECAR did not show any differences over 7 days, suggesting that TDP-43 A315T expression damages mitochondria while leaving glycolysis unaffected (**Fig 1j**).

### Expression of TDP-43 A315T does not result in nuclear TDP-43 loss of function

A pathological hallmark of ALS is the accumulation of cytoplasmic TDP-43 aggregates accompanied by depletion of nuclear TDP-43^1^. To investigate nuclear localization of mutant TDP-43 we monitored its localization though the Dendra2 tag using both nuclear fractionation by WB and confocal microscopy (**Fig. 2a, d**). WB analysis did not reveal differences in the nuclear localization of TDP-43 A315T compared to wild-type TDP-43 (69 kDa band, **Fig. 2b**). Similarly, the nuclear levels of endogenous untagged TDP-43, which is wild type in both cell lines, were not affected by the presence of the A315T mutation (43 kDa band, **Fig. 2c**). Consistent with these findings, quantitative confocal microscopy revealed no differences in nuclear Dendra2 fluorescence between TDP-43 WT and TDP-43 A315T cortical neurons, indicating that the A315T mutation does not alter basal nuclear localization of TDP-43 (**Fig. 2d, e**).

**Figure 2.**
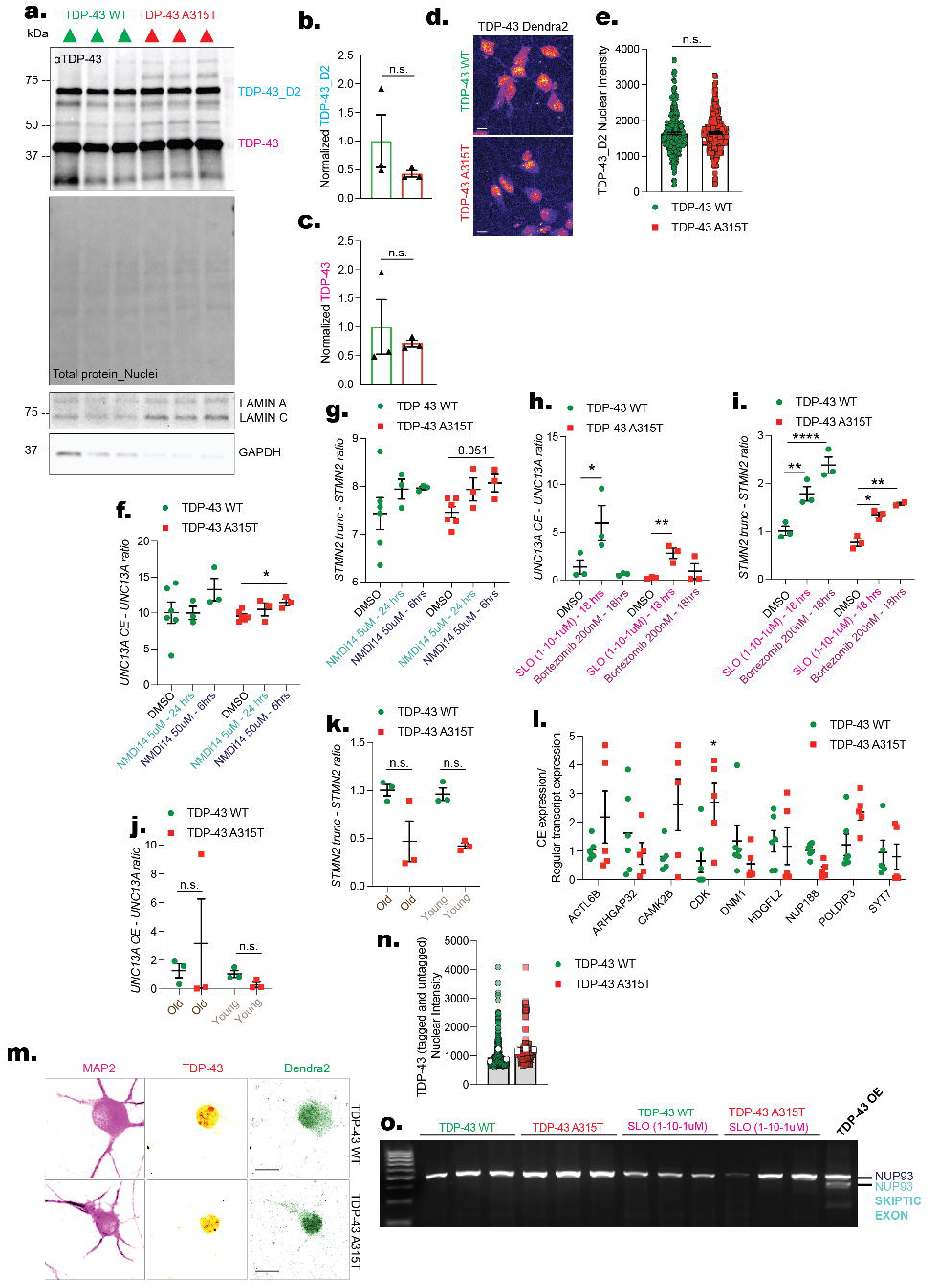
TDP-43 A315T neurons retain nuclear TDP-43 at baseline but display selective loss of nuclear function upon stress. (a) Representative immunoblot of nuclear fractions from TDP-43 WT and TDP-43 A315T I3 neurons probed with anti-TDP-43 antibody (αTDP-43), LAMIN A, LAMIN C, and GAPDH. Total protein is revealed through Ponceau S. (b–c) Quantification of normalized TDP-43_Dendra2 (b) and total TDP-43 (c) band intensity in nuclear fractions. Data presented as mean ± SEM, n = 3 independent replicates. Not significant, Student t-test. (d) Representative confocal immunofluorescence images of Dendra2 signal (color range reporting for Dendra2 signal intensity) in TDP-43 WT (top) and TDP-43 A315T (bottom) I3 neurons. (e) Quantification of TDP-43_Dendra2 nuclear intensity from single-cell imaging in TDP-43 WT (green) and TDP-43 A315T (red) neurons. Data presented as mean ± SEM, n = 3 independent replicates, m> 100 cells per replicate. Not significant, Student t-test. (f-g) UNC13A CE/UNC13A (f) and STMN2 truncated/STMN2 (g) ratios in TDP-43 WT and TDP-43 A315T neurons treated with the NMD inhibitor NMDi14 (5 µM for 24 hours or 50 µM) for 48 hours) or DMSO vehicle. Data shown as mean ± SEM, n = 3–6 independent replicates. *p<0.05, Student t-test. (h–i) UNC13A CE/UNC13A (h) and STMN2 truncated/STMN2 (i) ratios in TDP-43 WT and TDP-43 A315T neurons treated with SLO (1–10 µM) or with Bortezomib (200 nM) for 18 hours. Data shown as mean ± SEM, n = 3 independent replicates. Data shown as mean ± SEM, n = 3–6 independent replicates. *p<0.05, **p<0.01, ***p<0.001, ****p<0.0001 One-Way ANOVA. (j–k) UNC13A CE/UNC13A (j) and STMN2 truncated/STMN2 (k) ratios in old (DIV35) versus young (DIV21) TDP-43 WT and TDP-43 A315T neurons. Data shown as mean ± SEM. n = 3–6 independent replicates. Not significant, One-Way ANOVA. (l) Quantification of cryptic exon to regular transcript expression ratios across a panel of TDP-43 target genes (ACTB, ARHGAP32, CAMK2B, CDK, DNM1, HDGFL2, NUP188, POLDIP3, SYT7) in TDP-43 WT and TDP-43 A315T neurons (DIV21), relative to the corresponding regular transcript. Each dot represents an independent biological replicate. Not significant, One-Way ANOVA. (m) Representative confocal images of single TDP-43 WT and TDP-43 A315T neurons immunostained for endogenous TDP-43 (αTDP-43; yellow) and the dendritic marker MAP2 (magenta) and showing Dendra2 in green. Scale Bar = 10μm (n) Quantification of nuclear TDP-43 intensity in TDP-43 WT and TDP-43 A315T neurons. Each dot represents one cell. Data presented as mean ± SEM, n = 3 independent replicates, m> 100 cells per replicate. Not significant, Student t-test. (o) Agarose gel electrophoresis of RT-PCR products using NUP93 exon-flanking primers that detect both the canonical NUP93 band and a lower-molecular-weight NUP93 skiptic exon product.

Next, we investigated whether TDP-43 A315T can affect wild-type TDP-43 nuclear function. Nuclear loss of TDP-43 disrupts its RNA-binding function, resulting in aberrant RNA processing, including cryptic exon incorporation into transcripts such as STMN2^9^ and UNC13A^8^, two well-established downstream markers of TDP-43 loss of function. We therefore asked whether endogenous expression of the A315T mutation was sufficient to induce nuclear depletion and impair TDP-43-mediated RNA processing. We quantified both the physiological and cryptic exon-containing transcripts of UNC13A and STMN2. Cryptic exon inclusion was expressed as the ratio of aberrant to canonical transcripts. At baseline, TDP-43 WT and TDP-43 A315T neurons exhibited comparable levels of UNC13A and STMN2 cryptic exon inclusion, indicating preserved TDP-43 splicing activity despite the presence of the A315T mutation (**Fig. 2f, g**). To determine whether rapid degradation of aberrant transcripts masked subtle defects in TDP-43 splicing, we inhibited non-sense mediated decay (NMD) using NMDi14^30,31^. Blocking NMD stabilizes cryptic exon-containing transcripts, allowing their accumulation and more accurate quantification. We used two doses and treatment duration to cover a broader spectrum of degradation kinetics. Interestingly, both lines show a mild increase in aberrant transcripts expression compared to untreated controls looking at both UNC13A and STMN2 (**Fig. 2f, g**). This trend reached significance only in case of UNC13A-CE in TDP-43 A315T mutant lines when treatment concentration was increased by 10 fold and kept for 6 hours (**Fig. 2f**). Interestingly, NMDi14 treatment also increased the expression of total UNC13A and STMN2 in both lines (**Fig. S3a**). Surprisingly also GAPDH transcripts were found upregulated in presence of NMD blockage (**Fig. 3a**), and was thus not used as normalizer. To further investigate TDP-43 loss of function in our lines, we used SLO and Bortezomib to rapidly induce senescence and block proteasome function, respectively, thereby artificially inducing TDP-43 nuclear translocation^32,33^. Notably both cell lines responded to SLO with higher rate of aberrant transcript inclusion in both UNC13A and STMN2 genes (**Fig. 2h, i**), whereas bortezomib induced in both lines the aberrant truncation of STMN2, without affecting the rate of UNC13A CE inclusion events (**Fig. 2h, i**). Because aging is the strongest risk factor for ALS and has been associated with progressive TDP-43 dysfunction, we asked whether prolonged neuronal maturation would unmask defects in TDP-43 mediated RNA processing. We extended the culture period from 21 to 35 DIV and re-evaluated cryptic exon inclusion in UNC13A and STMN2. Prolonged culture did not increase cryptic exon incorporation, and TDP-43 WT and TDP-43 A315T neurons remained indistinguishable at both time points, indicating preservation of basal TDP-43 splicing activity despite neuronal aging in vitro (**Fig. 2j, k**). While the complete repertoire of transcripts processed by TDP-43 is unknown, we expanded our analysis to include ACTL6B, ARHGAP32, CAMK2B, CDK, DNM1, HDGFL2, NUP188, POLDIP3, and SYT7, which are reported targets of TDP-43^34^. Quantitative PCR analysis did not reveal robust and significant increase of these transcripts in the mutant lines (**Fig. 2l**), with the exception of CDK, whose aberrant variant was three times higher in mutant lines compared to wild type.

**Figure 3.**
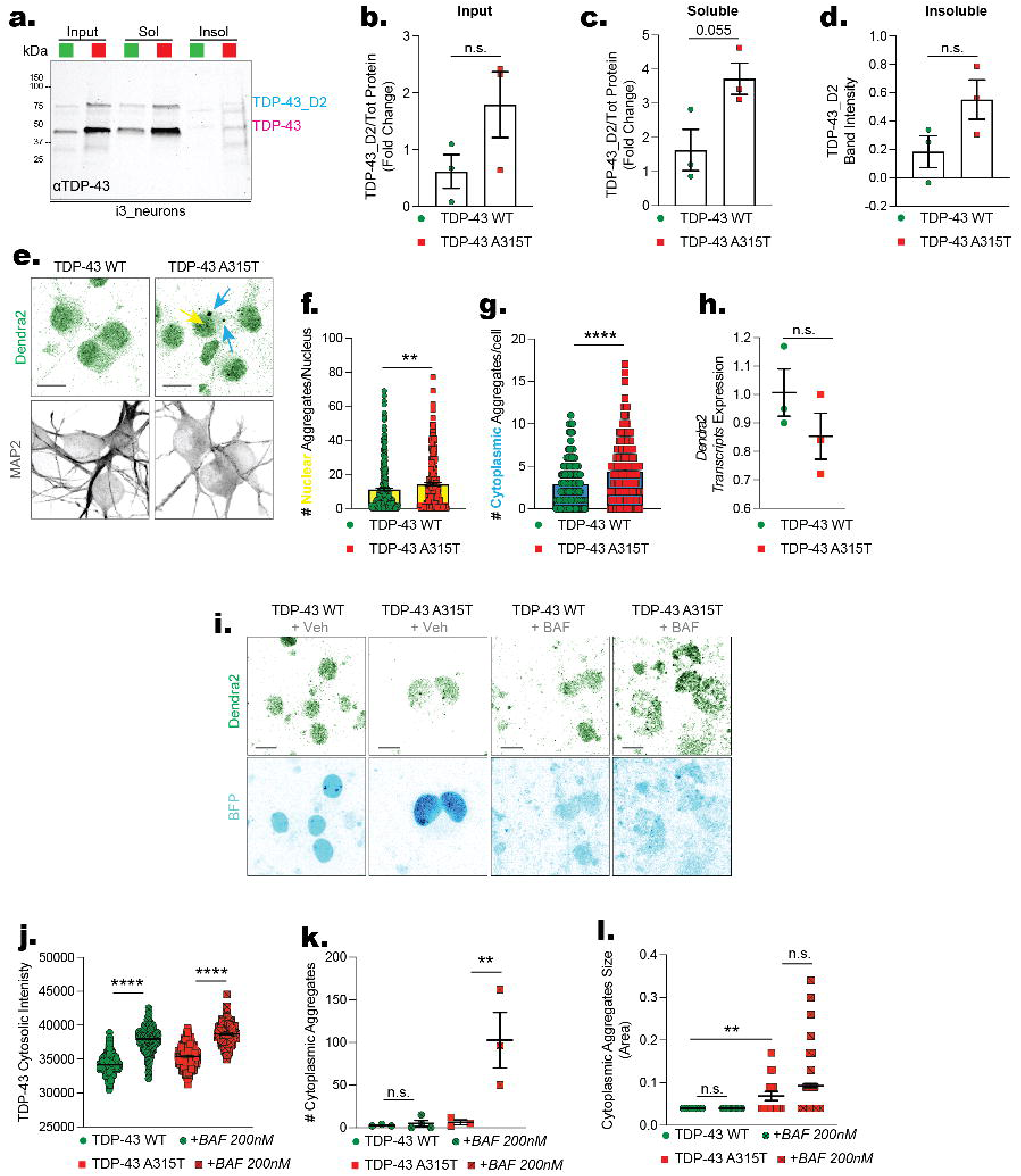
TDP-43 A315T neurons accumulate cytoplasmic TDP-43 aggregates that are modulated by autophagy. (a) Representative immunoblots of input, soluble (Sol), and insoluble (Insol) fractions from TDP-43 WT and TDP-43 A315T neurons probed with anti-TDP-43 antibody (αTDP-43) (b–d) Quantification of TDP-43_Dendra2 band intensity in input (b), soluble (c), and insoluble (d) fractions, normalized to Ponceau S total protein staining. Data presented as mean ± SEM, n = 3–4 independent experiments. p = 0.055 for soluble fraction (c). (e) Representative confocal fluorescence images of TDP-43 WT and TDP-43 A315T neurons stained for Dendra2 (TDP-43; green) and MAP2 (gray). Yellow arrows indicate nuclear TDP-43 aggregates; cyan arrows highlight cytoplasmic aggregates in A315T neurons. Scale Bar = 10µm (f-g) Quantification of nuclear (f) or cytoplasmic (g) TDP-43_Dendra2 aggregates per nucleus/cell in TDP-43 WT (green) and TDP-43 A315T (red) neurons. Data presented as mean ± SEM, n = 3 independent replicates, m> 100 cells per replicate. **p<0.01, ****p<0.0001 Student t-test. (h) RT-qPCR quantification of **TDP-43 Dendra2** transcript expression in TDP-43 WT and TDP- 43 A315T neurons, further confirming downregulation at the transcript level. Data presented as mean ± SEM. Non significant. (i) Representative confocal fluorescence images of TDP-43 WT and TDP-43 A315T neurons treated with vehicle (Veh) or BAF (200 nM), showing Dendra2 (TDP-43; green) and BFP (blue). Scale Bar = 10µm (j) Quantification of **total cytoplasmic TDP-43 fluorescence** intensity per cell in TDP-43 WT and TDP-43 A315T neurons treated with vehicle (Veh) or BAF (200 nM). Data presented as mean ± SEM, n = 3 independent replicates, m> 100 cells per replicate. ****p<0.0001 One-Way ANOVA. (k) Quantification of **cytoplasmic aggregate size** (arbitrary area units) in TDP-43 WT and TDP-43 A315T neurons treated with vehicle (Veh) or BAF (200 nM). Data presented as mean ± SEM, n = 3 independent replicates, m> 100 cells per replicate. **p<0.01, One-Way ANOVA. **p<0.01, Student t-test. (l) Quantification of **cytoplasmic TDP-43_Dendra2 aggregates per field** in TDP-43 WT and TDP-43 A315T neurons treated with vehicle (Veh) or BAF (200 nM). Data presented as mean ± SEM, n = 3 independent replicates, m> 100 cells per replicate. **p<0.01, One-Way ANOVA.

We then asked whether the total levels of TDP-43 (modified allele + unmodified) were affected through total TDP-43 staining followed by confocal microscopy and we observed a slight increase in the average TDP-43 fluorescence in mutant compared to wild type lines (**Fig. 2m, n**). Functionally, increased nuclear TDP-43 levels can lead to aberrant exon skipping^35,36^, an event that can also influence cell health and function. We thus performed end-point PCR using specific primers to measure the skiptic events in the NUP93 mRNA. We did not observe the appearance of skiptic exons in either line (**Fig. 2o**). The specificity of this assay was confirmed using complementary gain- and loss-of-function controls. SLO-mediated depletion of nuclear TDP-43 produced the expected reduction in the canonical NUP93 transcript, whereas TDP-43 overexpression in HEK293 cells induced robust skiptic exon formation (**Fig. 2o**).

In conclusion, these results support a model in which expression of mutant TDP-43 in mature neurons causes mitochondrial dysfunction, reduced neurite outgrowth, and aberrant electrical activity. Yet, these effects cannot be explained by canonical TDP-43 dysfunction as indicated by the absence of aberrant exon inclusion or skipping.

### TDP-43 A315T aggregation in the cytoplasm of cortical neurons is regulated by autophagy activity

Because the A315T mutation lies within the aggregation-prone C-terminal low-complexity domain of TDP-43, we next examined whether endogenous expression of the mutant TDP-43 promoted spontaneous aggregation in human cortical neurons. Whole cell lysates were separated into RIPA-soluble and insoluble fractions to quantify soluble and aggregated TDP-43 species (**Fig 3a**). WB analysis showed a clear trend with higher levels of TDP-43 A315T in the both soluble and insoluble fractions compared to the wild type counterpart (69kDa band) (**Fig. 3a-d**). Pathological TDP-43 aggregates are usually found in the cytosol, which cannot be discerned by biochemical fractionation. We therefore used high content imaging to differentiate between nuclear and cytosolic aggregates at the single cell-level (**Fig 3e-g**). Strikingly, we observed a robust increase in the average number of TDP-43 A315T Dendra2 nuclear aggregates (yellow arrow, **Fig 3e, f**) compared to wild type TDP-43 Dendra2. We then also counted the aggregates present in the cytoplasm of the cortical neurons and again found an increase in the number of TDP-43 A315T Dendra2 aggregates per cell compared to wild type TDP-43 Dendra2 (blue arrow, **Fig 3e, g**). To assess whether the increase in aggregation was attributable to increased mRNA expression, we measured the expression levels of the edited allele by qPCR using primers specific for Dendra2. We found that TDP-43 Dendra2 expression was comparable between the two lines (**Fig 3 h**), suggesting that differences in protein degradation, together with the mutation itself, rather than differences in protein expression, account for their differential aggregation propensity.

One of the possible routes of TDP-43 degradation in the cytoplasm is autophagy which can also eliminate protein aggregates. Conversely, accumulation of aggregates overtime can significantly affect autophagy function eventually leading to its blockage^37^. Autophagy levels are evaluated by LC3I and LC3II levels. Increased levels of LC3-II signify autophagy blockage when not paralleled by an increase in LC3-I^38^. Using lysates from our cortical neuron cell lines, we detected a robust increase in LC3-II and the LC3-II/LC3-I ratio (**Fig S4a-d**), indicating autophagy involvement in neurons endogenously expressing TDP-43 A315T. To discriminate between autophagy activation or blockage in presence of TDP-43 A315T mutation, we inhibit the last step of autophagy which is the fusion between lysosomes and autophagosomes with bafilomycin (**Fig 3i-l**). Through confocal microscopy we measure TDP-43 cytoplasmic mislocalization and the formation of cytosolic aggregates. Notably, we observed a significant increase in TDP-43 Dendra2 localization in the cytoplasm in both cell lines upon autophagy inhibition, indicating autophagy involvement in the degradation of both wild type and mutant cytosolic species (**Fig 3j**). Strikingly, autophagy inhibition increased the formation of TDP-43 aggregates exclusively in the presence of the A315T mutation (**Fig 3k**) without affecting their size (**Fig 3l**).

### TDP-43 A315T enhances stress granule formation and recruitment of TDP-43 under oxidative stress

Liquid-liquid phase separation of protein is the physical phenomenon underlying stress granule formation process also driven by specific cytosolic RNAs^31^. SGs have been postulated to act as seeding sites for aggregates formation. We reasoned that SGs formation and content might be influenced by the A315T mutation, given that the location of the mutation is at the C-terminus domain responsible for TDP-43 phase separation^39^. Notably, literature around this specific mutation and its interaction with SGs is very limited.

We used sodium arsenite to induce SGs formation through oxidative stress and marked SGs through G3BP1 the most abundant SG protein. In unchallenged cells, we observed spontaneous formation of SGs in the cytoplasm of cells at a higher rate in neurons harboring the A315T mutation, as shown by the increased number of G3BP1^+^ SGs (**Fig 4a, b**). While the size of the SGs at baseline was unchanged between genotypes (**Fig 4a, c**), TDP-43 A315T, measured through Dendra2 intensity, was more prone to accumulate in SGs than the wild-type counterpart (**Fig 4a, e**). Upon sodium arsenite stimulation, we observed a strong increase in SGs number and size in both lines compared to their respective controls (**Fig 4a, b**). Notably, TDP-43 A315T neurons upon arsenite treatment form bigger SGs than the ones formed in the cytoplasm of the control neurons (**Fig 4a, c**). Interestingly, the number of TDP-43 aggregates increased in both cell lines in response to oxidative stress (**Fig 4a, d**). Lastly, as expected, sodium arsenite treatment led to higher localization of TDP-43 A315T Dendra2 protein within the stress granules compared to the untreated group and to the isogenic control line (**Fig 4a, e**).

**Figure 4.**
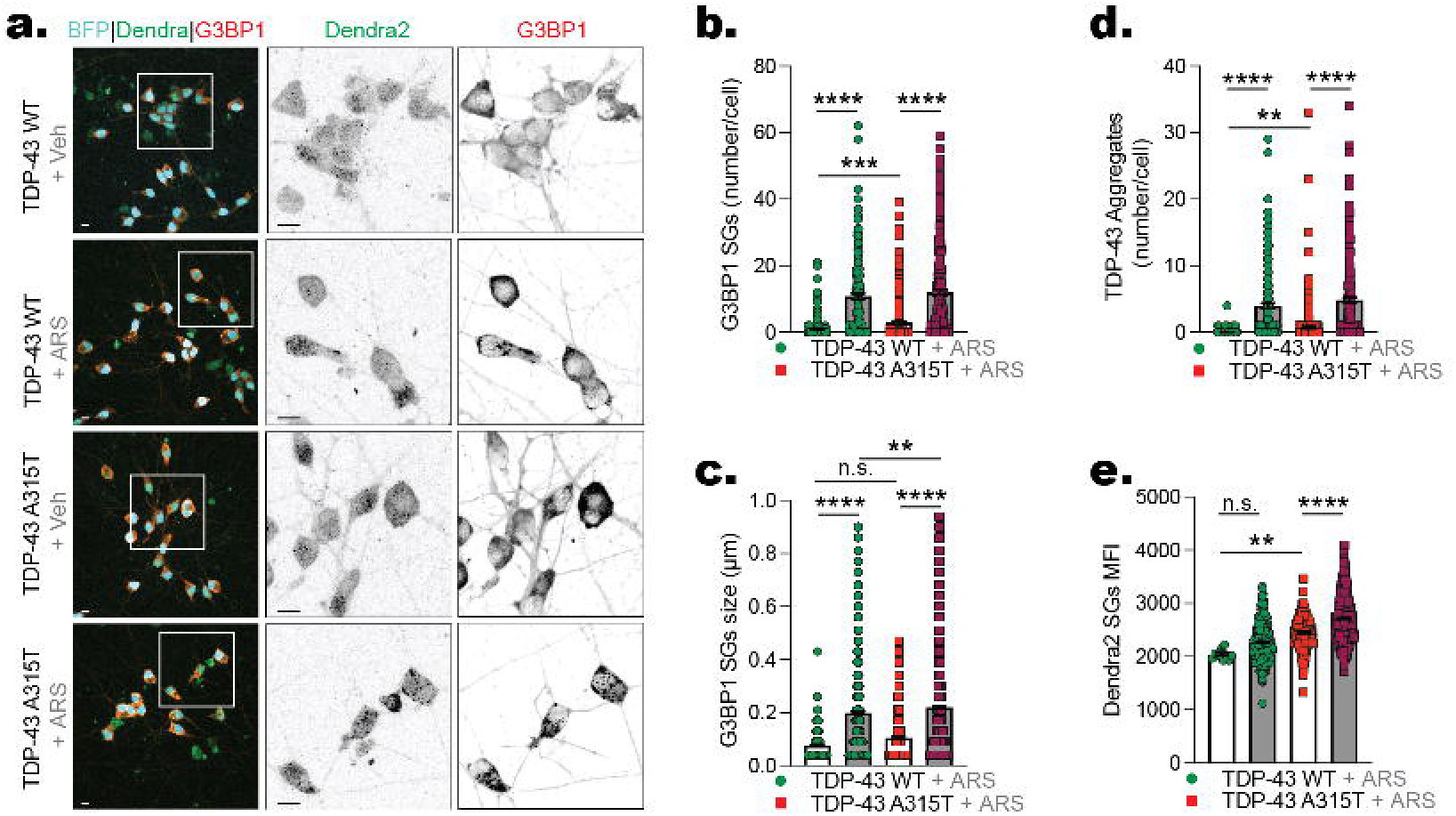
TDP-43 A315T localizes within stress granules following arsenite-induced oxidative stress. (a) Representative confocal fluorescence images of TDP-43 WT and TDP-43 A315T I3 neurons treated with vehicle (Veh) or sodium arsenite (ARS) and immunostained for G3BP1 (red/gray); BFP (nuclear marker; cyan), Dendra2-TDP-43 (green). Scale Bar = 10µm (b) Quantification of the number of G3BP1-positive stress granules per cell across conditions. Data presented as mean ± SEM, n = 3 independent replicates, m> 100 cells per replicate. ***p<0.001, ****p<0.0001 One-Way ANOVA. (c) Quantification of the size of G3BP1-positive stress granules per cell across conditions. Data presented as mean ± SEM, n = 3 independent replicates, m> 100 cells per replicate. ***p<0.001, ****p<0.0001 One-Way ANOVA. (d) Quantification of TDP-43-positive aggregates per cell across conditions. Data presented as mean ± SEM, n = 3 independent replicates, m> 100 cells per replicate. ***p<0.001, ****p<0.0001 One-Way ANOVA. **p<0.01, Student t-test. (e) Quantification of Dendra2-positive SG mean fluorescence intensity (MFI) per cell. Data presented as mean ± SEM, n = 3 independent replicates, m> 100 cells per replicate. **p<0.01, ***p<0.001, One-Way ANOVA.

In conclusion, the expression of the aggregation-prone TDP-43 A315T was sufficient to stimulate SGs formation while only following acute oxidative stress with sodium arsenite, it caused an increase in TDP-43 aggregates and an increase in its own localization to SGs. Thus, our data support a model in which mutant TDP-43 can influence other phase-separated protein aggregates and its own phase separation in response to oxidative stress.

### Clip34 suppresses TDP-43 A315T aggregation and alters its partitioning into stress granules

TDP-43 aggregation is a pathological hallmark of ALS and several other neurodegenerative and muscular diseases, making it an attractive therapeutic target. We reasoned that our cortical neuronal model could be successfully used as a therapeutic discovery platform. We thus decided to focus on Clip34, an RNA aptamer previously shown to reduce TDP-43 aggregation through binding to its RNA-recognition motif (RRM)^40,41^. Notably, its activity in the context of the ALS-associated A315T mutation has never been examined. We thus delivered Clip34 in mature cortical neurons expressing either wild-type or A315T alleles, with or without sodium arsenite, and assess TDP-43 aggregation and SGs parameters. Strikingly, at baseline, Clip34 significantly reduced the number of cytoplasmic TDP-43 aggregates in both WT and A315T neurons (**Fig. 5a,b**). As expected, arsenite-induced oxidative stress markedly increased TDP-43 aggregation; this response was significantly attenuated by Clip34 treatment in both lines (**Fig 5a, b**). These findings demonstrate that Clip34 suppresses TDP-43 aggregation under both basal and stress conditions and establish the aggregation phenotype in our model as amenable to therapeutic modulation.

**Figure 5.**
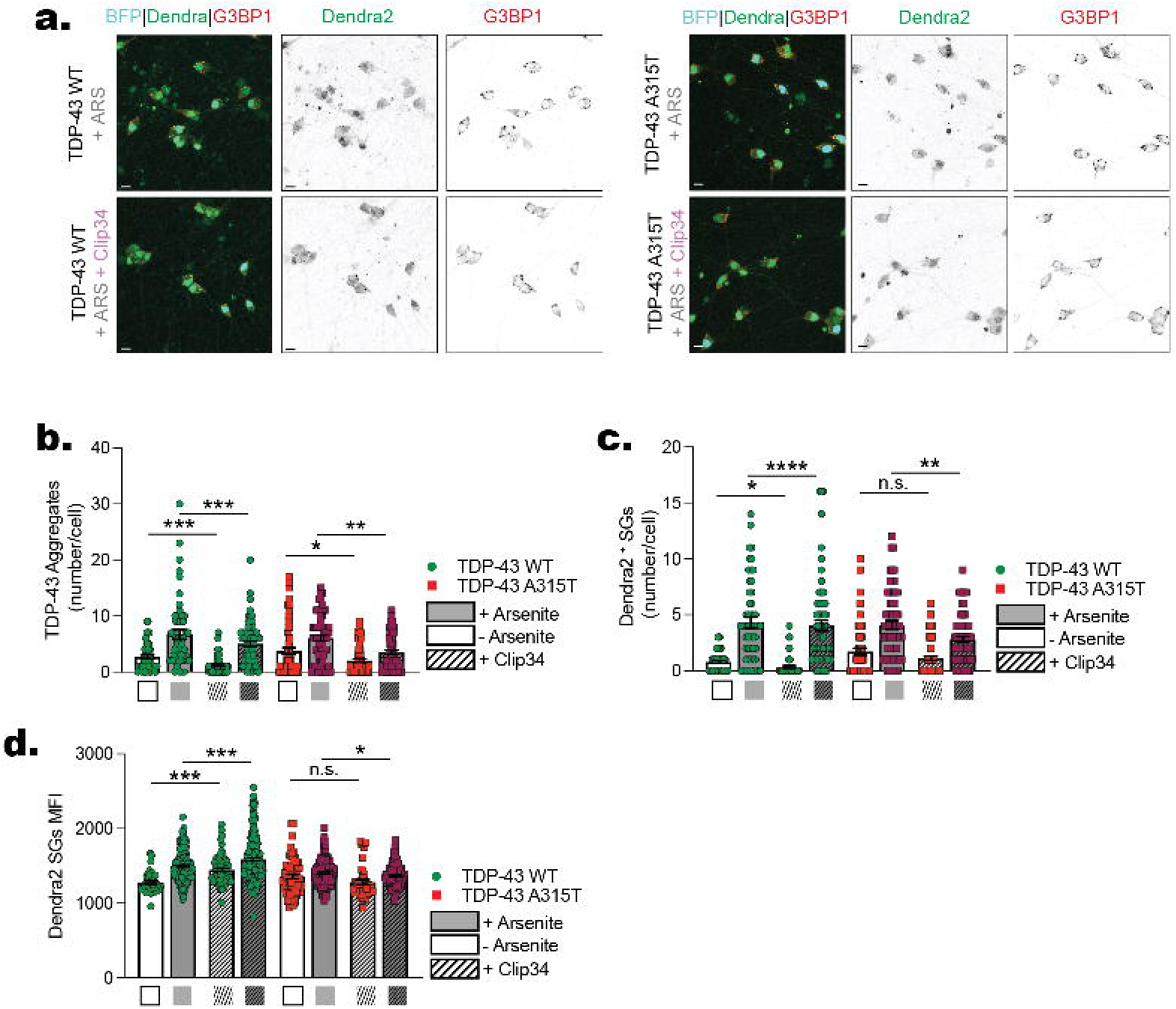
The oligonucleotide Clip34 reduces TDP-43 aggregation without affecting stress granule dynamics. (a) Representative confocal fluorescence images of TDP-43 WT and TDP-43 A315T I3 neurons treated with ARS ± Clip34, immunostained for G3BP1 (red/gray); BFP (nuclear marker; cyan), Dendra2-TDP-43 (green). For each genotype, ARS alone and ARS + Clip34 conditions are shown. Scale Bar = 10µm (b) Quantification of **number of TDP-43_Dendra2-positive aggregates per cell** in TDP-43 WT and TDP-43 A315T I3 neurons treated with ARS ± Clip34. Data presented as mean ± SEM, n = 3 independent replicates, m> 100 cells per replicate. **p<0.01, ***p<0.001 One-Way ANOVA. *p<0.05, ***p<0.001 Student T-test. (c) Quantification of **Dendra2-positive stress granule number per cell** in TDP-43 WT and TDP-43 A315T I3 neurons treated with ARS ± Clip34. Data presented as mean ± SEM, n = 3 independent replicates, m> 100 cells per replicate. *p<0.05, **p<0.01, ***p<0.001, ****p<0.0001 One-Way ANOVA. *p<0.05, **p<0.01 Student T-test. (d) Quantification of **Dendra2 mean fluorescence intensity per SGs** in TDP-43 WT and TDP-43 A315T I3 neurons treated with ARS ± Clip34. Data presented as mean ± SEM, n = 3 independent replicates, m> 100 cells per replicate. *p<0.05, **p<0.01, ***p<0.001, ****p<0.0001 One-Way ANOVA. *p<0.05, **p<0.01 Student T-test.

Through G3BP1 staining we analyzed the number of SGs per cell positive to Dendra2, thus containing either TDP-43 wild type Dendra2 or TDP-43 A315T Dendra2 (**Fig 5a, c**). We again observed, at baseline, an increase in the number of SGs positive for Dendra2 in neurons expressing mutant TDP-43 (**Fig 5a, c**). Clip34 reduced this phenotype, although the effect did not reach statistical significance (**Fig a, c**). Upon oxidative stress both lines responded by increasing the number of SGs containing TDP-43. Clip34 led to a reduction of ∼50% in the formation of SGs containing TDP-43 in both lines, although it did not completely avert SGs formation (**Fig 5 a, c**). Finally, we quantified the amount of Dendra2-tagged TDP-43 within individual SGs and observed a mutation-dependent response to Clip34. In the wild type line, Clip34 led to an increase in the TDP-43 content of SGs both at baseline and under oxidative stress (**Fig 5 a, d**). In contrast, cortical neurons expressing TDP-43 A315T present a reduction at baseline which did not reach statistical significance and instead a significant reduction in the TDP-43 content in presence of oxidative stress, possibly suggesting that the mutation leads to a different interaction with the Clip34 molecule (**Fig 5 a, d**). These findings suggest that the A315T mutation alters the response of TDP-43 to Clip34 and further demonstrate the ability of this model to resolve mutation-specific responses to candidate therapeutic interventions.

## Discussion

In this study, we established a human cell–based model using the ALS-linked A315T mutation as a genetic entry point for investigating TDP-43 dysfunction. A315T is considered a particularly pathogenic TDP-43 variant and has been associated in animal models with neurodegeneration, neuroinflammation, and widespread molecular abnormalities, partly through its effects on TDP-43 solubility. To preserve physiological expression and genetic context, we used CRISPR–Cas9 genome editing to introduce the A315T mutation in heterozygosis into exon 6 of the endogenous TARDBP locus, recapitulating the genotype found in patients. We also fused TDP-43 to the photoconvertible fluorescent protein Dendra2, enabling spatial and temporal tracking of the endogenous protein. Importantly, a previous study used a similar Dendra2-based approach to measure TDP-43 turnover in neurons and reported that the tag did not detectably alter TDP-43 half-life, including in the presence of the A315T mutation^42^. In this study TDP-43 A315T had a significant lower half-time compared to wild type and that autophagy modulation influences the stability of both proteins. Although here we did not exploit Dendra2’s green-to-red photoconversion to measure protein half-life this feature positions the tool for future studies of TDP-43 stability and turnover.

We used iPSCs for their versatility and capacity to generate diverse cell types and more complex systems. Inducible cortical neurons offer a tractable, monoculture system that is easy to manipulate and yields straightforward, interpretable results. Strikingly, the A315T line diverged from wild type within days: neurite length, firing activity, and oxygen consumption — measured by two independent methods — were all reduced. These three phenotypes points toward a single explanation, mitochondrial dysfunction, since defective mitochondria lower ATP production and thereby impair neuronal activity and synaptic transmission. Since we measured oxygen consumption in Brainphys medium with 2.5 mM glucose which matches the physiological brain glucose levels, we are confident that the differences observed are the result of an intrinsic metabolic defects rather than the consequence of a non physiological glucose concentration. This interpretation is supported by GSEA of our RNAseq data, which revealed pronounced dysregulation of mitochondrial pathways. Although there is evidence describing the role of TDP-43 in mitochondria^14,17–20,43^, a full picture of the physiological role of TDP-43 within these organelles remains unresolved and warrants direct investigation in the context of this mutation. However, regardless of mechanism, the early onset of these phenotypes makes this model attractive for therapeutic screening in ALS.

We next asked whether the mutation, which lies outside the RNA Recognition Motif (RRM) responsible for TDP-43’s role in RNA maturation, disrupts its nuclear function. A substantial body of recent literature has shown that TDP-43 loss of function drives aberrant retention of introns as cryptic exons, which are then degraded via nonsense-mediated decay or related pathways^7–10,12^. Yet our A315T line showed no difference from wild type in the maturation of STMN2 and UNC13A, the transcripts most strongly affected by TDP-43 pathology^8,9^ in ALS and related dementias, at either early or late time points. We confirmed that our model is capable of generating cryptic exons when TDP-43 nuclear localization is chemically disrupted. However, under baseline conditions, the two lines were indistinguishable, and inhibition of nonsense-mediated decay revealed only a modest difference in RNA turnover in the mutant line. These findings were not unexpected, as TDP-43 loss of function in neuronal cultures has primarily been demonstrated following TDP-43 depletion or forced nuclear mislocalization, neither of which occurs in our model under baseline conditions. The absence of nuclear mislocalization is consistent with the literature and confirms that our data align with the current understanding of TDP-43’s role in RNA maturation. Combined with the mutation’s clear effects on TDP-43 solubility and RNA binding, this gives us confidence that our model faithfully captures A315T behavior.

Prior work *in vitro*, in cell culture, and in animal models has shown that the A315T mutation markedly reduces TDP-43 solubility, driving faster aggregation into inclusions that resist clearance^15,26,44^. Neurons are typically vulnerable to aggregate formation, in part because, as post-mitotic cells, they cannot dilute aggregates through division, and in our model, aggregates were already detectable in both the nucleus and cytoplasm by day 21 *in vitro*. In contrast to the nucleus, cytoplasmic aggregate accumulation tracked with divergent autophagic activity between the mutant and isogenic control lines: baseline LC3-I-to-LC3-II conversion, which drives autophagosome maturation, already differed between lines, and blocking autophagic flux sharply increased cytoplasmic aggregate accumulation. Together, these results show that although autophagy actively degrades mutant TDP-43, its capacity is insufficient to fully clear the mutant protein and to prevent aggregate accumulation at baseline

Aggregates are thought to exert toxicity in part by sequestering physiological proteins, disrupting the formation of LLPS-based structures impairing their function^45^. We therefore examined stress granules (SG) formation, since SGs likewise assemble through liquid-liquid phase separation mediated by low-complexity domains. In unchallenged neurons, spontaneous SG formation and TDP-43 content within SGs were both elevated presence of the mutation, a pattern consistent with faster TDP-43 condensation acting as a nucleating seed for SG assembly. Oxidative stress induced by sodium arsenite acutely amplified SG formation and TDP-43 aggregation in both lines, but the mutant line showed a higher increase in SG size and TDP-43 content, indicating that A315T sensitizes this pathway to stress. To probe the nature of these aggregates, we used Clip34, a small oligonucleotide that is recognized by the TDP-43 RRM and reduces TDP-43 aggregation propensity^40,41^; while Clip34 was previously shown to decrease oligomerization of wild-type TDP-43^41^, it had never been tested against aggregates formed by mutant TDP-43. We show here that TDP-43 A315T aggregates are also sensitive to Clip34: aggregation was significantly reduced both at baseline and under oxidative stress. Clip34 also reduced SG formation under oxidative stress in the mutant line and lowered TDP-43 content within SGs, further linking TDP-43 aggregation to SG dynamics. These data corroborate and reinforce recent literature supporting Clip34 as avenue to ameliorates TDP-43 aggregation.

In conclusion, this mutant heterozygous model robustly recapitulates core TDP-43 pathological features — neurodegeneration, metabolic imbalance, and protein aggregation — each easily monitored over time, making it well suited for therapeutic development without reliance on large animal cohorts and in settings closely recapitulating the human disease. Extending this platform to more complex systems, such as 3D organoid cultures, will further enrich the toolbox available for studying TDP-43 pathology. We believe this model fills an important gap in the field, by providing a physiologically relevant cellular platform for investigating TDP-43-associated neuronal dysfunction.

### Limitation of the Study

The goal of the present study is to showcase a new powerful tool to study TDP-43 dynamics and associated pathological mechanisms in human setting. Importantly, our system closely resembles patient case as it presents the mutation in heterozygosis. However, from a functional standpoint we cannot distinguish the contribution of mutant TDP-43 from that of the WT untagged protein. Secondly, the study is carried out in two isogenic cell lines. Future studies including patients derived cell lines are warranted in order to expand and strengthen these results. Last, the linker-mediated Dendra2 fusion to the TDP-43 protein might bring subtle changes to the biophysical properties of the protein. While we cannot rule out this possibility, previous studies used TDP-43-Dendra2 showing no alteration of major behavioral features of the protein.

## Author Contribution

**SA and MEC** performed the experiments, collected and analyzed the data, prepared the figures, and contributed to manuscript revision. **IMS** provided technical guidance on stem cell maintenance and quality control. **MM** supervised the generation and characterization of i³Neurons. **NK** performed and interpreted the skiptic exon analyses. **AG** provided Clip34 and advised on the Clip34-related studies. **AP and JS** provided guidance and supervision for the CRISPR-Cas9 editing experiments. **VC and MEC** designed and performed the CRISPR–Cas9 genome-editing experiments and validated the resulting cell lines. **VC and RC** performed the experiments to validate the excision of the cassette from the lines. **MB and MT** aided in interpreting the mitochondrial studies and provided technical and conceptual guidance. **MT and LG** contributed to study design, scientific supervision, data interpretation, and manuscript revision. **MEC and DT** conceived and designed the study, supervised the research, analyzed and interpreted the data, prepared the figures, and wrote and revised the manuscript. **DT, PP, and LG** acquired funding, and **DT** provided overall project administration. All authors reviewed and approved the final version of the manuscript.

## Acknowledgements

We would like to thank Drs. Jonathan Ling (Johns Hopkins) and Parker Sandal (Northwestern) for graciously providing samples from TDP-43 overexpressing HEK293 cells used in Figure 2; Dr Ratti for providing the parental iPS cell line, which underwent CRISPR-Cas9 Genome editing; Dr Tedesco for handling the shipment of the cells; Dr Goldsmith for providing the plasmids for the piggyback-based integration of the NGN2-BFP cassette for neuronal induction.

## Materials and Methods

### CRISPR-Cas9 Gene Editing

Isogenic human iPSC lines harboring the TDP-43 A315T mutation with an in-frame C-terminal Dendra2 tag were generated by CRISPR-Cas9-mediated homology-directed repair (HDR). Guide RNAs targeting TDP-43 exon 6 were designed to minimize off-target activity (**Table 1**) and cloned into a Cas9 expression vector pX335B_hCas9_2XhU6_guides (refer to Ran et al.^46^, for plasmid generation). A double-stranded HDR donor template encoding the GCG→ACG (Ala315Thr) substitution and the Dendra2 coding sequence was inserted in the pUC57 backbone and co-delivered with the plasmid expressing Cas9/sgRNAs into parental iPSCs (GAC2) by nucleofection. The parental cell line has been kindly donated by Prof Antonia Ratti, Auxologico, Milan to the Poletti Lab under an approved Material Transfer Agreement. The use of this line has been approved by Comitato etico 2015_03_31_07. Clonal selection was performed by single-cell plating, and individual clones were screened using PCR with specific primers (**Table 1**). Positive clones were checked by flow cytometry to confirm robust Dendra2 expression. Correct insertion of the mutation was also checked by Sanger Sequencing, and each cell lines genome was sequenced by whole genome sequencing. The Zeocin Selection cassette was then removed through Cre Recombinase treatment and excised clones were selected though flow cytometry. Successfully edited clones were confirmed by sequencing and retained for further analysis.

**Table 1.**
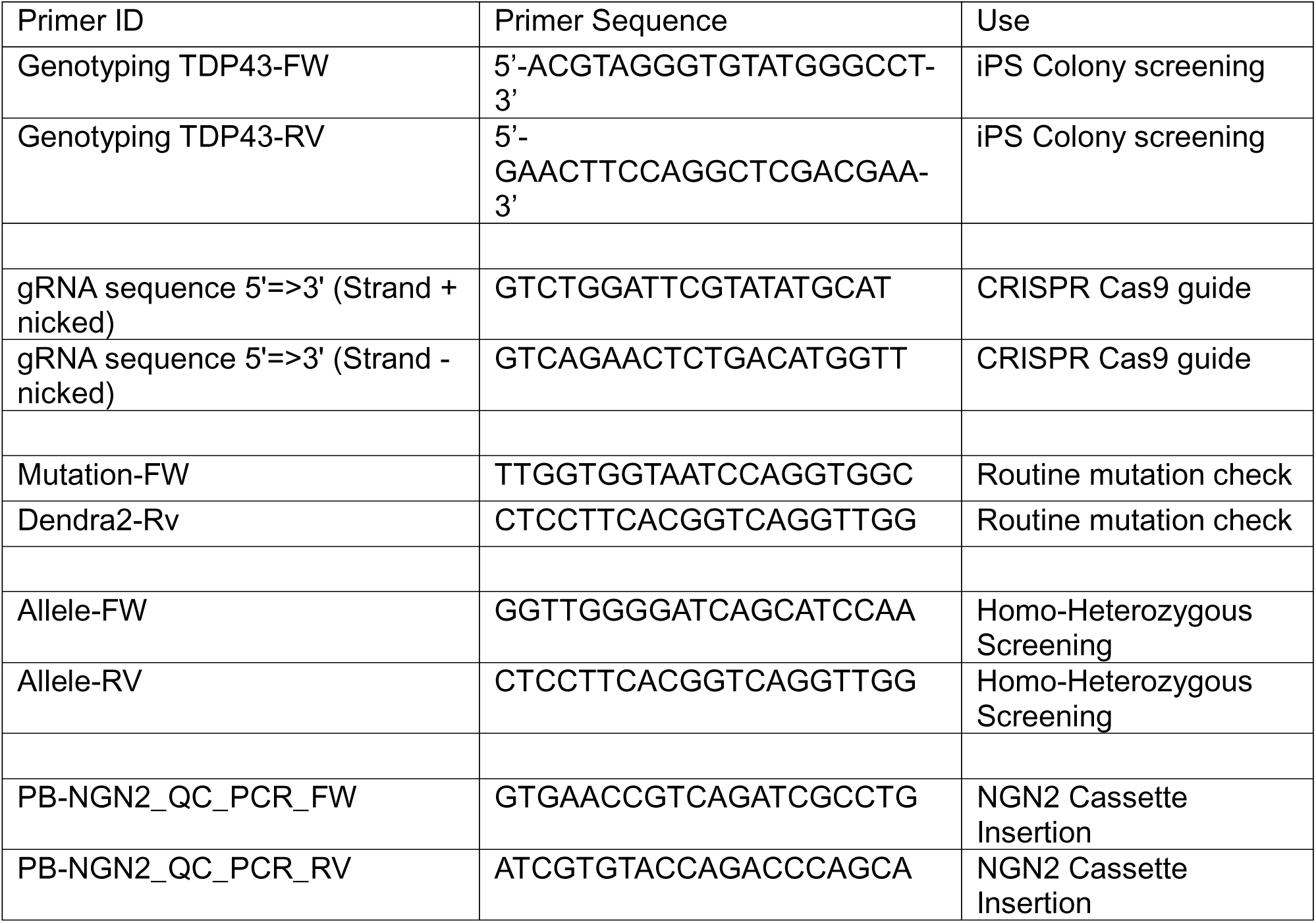
CRISPR-Cas9 Guides and PCR primers Sequences.

### Human Induced Pluripotent Stem Cell Culture

Parental and edited iPSC lines were maintained on Matrigel-coated (Corning) plates in mTeSR1 or StemFlex medium (STEMCELL Technologies) at 37°C in 5% CO2. Cells were passaged every 4–5 days using Accutase (STEMCELL Technologies) and plated at appropriate densities. All iPSC lines were routinely tested for mycoplasma contamination.

### Karyotyping

Chromosomal integrity of iPSC lines was assessed by standard G-band karyotyping. Cells at early passage were submitted to a certified cytogenetics laboratory for G-banding analysis. A minimum of 20 metaphases were analyzed per line. Results were compared to the karyotype of an unmodified iPSC line.

### DNA Sequencing

Genomic DNA was extracted from iPSC clones using the DNeasy Blood & Tissue Kit (Qiagen) following the manufacturer’s protocol. The region flanking the edited TDP-43 locus was PCR-amplified using primers specific to TDP-43 exon 6 and flanking intronic sequences (**Table 1**). PCR products were purified using the QIAquick PCR Purification Kit (Qiagen) and submitted for Sanger sequencing. Chromatograms were aligned to the reference human TARDBP sequence (GRCh38) to confirm successful introduction of the A315T substitution and Dendra2 tag.

### I3 iPSC Line Generation and Cortical Neuron Derivation and Maintenance

I3 neuron derivation was performed as described previously^47,48^. Briefly, iPSCs were transfected using 0.75 µg of PB-TO-hNGN2-puro-BFP donor plasmid (Plasmid #172115 Addgene) and 0.37 µg of EF1α-transposase plasmid (Plasmid #170824 Addgene) with 4 µL of Lipofectamine™ Stem in Opti-MEM. Stable integrants were selected with puromycin (1 µg/mL) for 5–7 days and maintained as I3-iPSC lines. Successful integration was confirmed by PCR amplification of the TRE-NGN2 insertion site (expected band: ∼320 bp (**Table 1**)) on a 1% agarose gel. BFP- and Dendra2-double-positive cells were sorted by FACS on day 2–3 to enrich for successfully edited cells.

For neuronal differentiation, I3-iPSCs were dissociated to single cells with Accutase, resuspended in induction media (N2 1%, NEAA 1%, DMEM/F12) supplemented with doxycycline (2 µg/mL) and ROCK inhibitor (10 µM Y-27632) and plated on Matrigel coated plates. Cells were freshly supplemented with doxycycline every 24 hours. After 3 days neurons were frozen or plated in maturation media (B27 2%, BDNF 10ng/mL, NT3 10ng/mL and laminin 1μg/mL in BrainPhys Neuronal Medium (STEMCELL Technologies)) on poly-L-ornithine(100μg/mL)/laminin(1μg/mL) coated plates. Neurons were matured up to 21 days and medium was half-changed every 3–4 days. Mature neurons were used for experiments between days 14 and 21 of differentiation unless otherwise specified.

### Flow Cytometry

For FACS-based sorting of iPSCs positive cells, cells were dissociated with Accutase on differentiation day 2–3, resuspended in FACS buffer (PBS + 2% FBS), and filtered through a 40 µm cell strainer. BFP and Dendra2 double-positive live cells were sorted using a Cytek Aurora CS spectral sorter. Sorting gates were established using single-color controls. Sorting experiments were performed in the Flow Cytometry Core Facility at Thomas Jefferson University.

### Alkaline Phosphatase

Pluripotency of iPSC colonies was evaluated by Alkaline Phosphatase (AP) staining using the Alkaline Phosphatase Detection Kit (Millipore, SCR004) following the manufacturer’s instructions. Briefly, iPSCs were fixed in 4% paraformaldehyde for 10 minutes at room temperature, washed with PBS, and incubated with the AP substrate for 15 minutes protected from light. Colonies were imaged under brightfield microscope equipped with color camera.

### Cell Treatment

Neurons were treated with the following compounds at the indicated concentrations and durations: Bafilomycin A1 (BAF, 200 nM); SLO (SBI-0206965 10 μM, Lopinavir 1 μM, O-151 1 μM); Bortezomib (200 nM); NMDi14 (5 or 150 µM); sodium arsenite (500 µM); Clip34 (500nM). All compounds were dissolved in DMSO (vehicle control) or water according to manufacturer instructions and diluted to working concentrations in neuronal maturation medium immediately prior to use.

### RNAseq

Total RNA was extracted from day 14–21 I3 neurons using TRIzol reagent (Thermo Fisher Scientific) following manufacturer instruction. RNA was quantified through Cytation 5 Agilent BioTek Cytation 5 Cell Imaging Multimode Reader. RNAseq was outsourced to Plasmidsaurus. Poly-A mRNA capture and reverse transcription for cDNA synthesis: oligo-dT primer to capture just the mRNA (contains sample barcode, UMI, and Read 1 sequences). Second strand synthesis and tagmentation: double stranded-cDNA is generated. Tagmentation creates fragments and incorporates Read 2 sequence. Illumina library amplification: adds unique dual indices (UDI = i5, i7) and P5/P7 sequences. Libraries were sequenced to a depth of ≥ 30 million reads per sample. Raw reads were aligned to the GRCh38 reference genome using STAR. Read counts were obtained with featureCounts (Subread package). Differential expression analysis was performed using DESeq2 in R, with a significance threshold of adjusted p-value < 0.05 and |log2FC| > 1. Gene Set Enrichment Analysis (GSEA) was performed using the fgsea package with Hallmark gene sets from MSigDB; Normalized Enrichment Scores (NES) and FDR-adjusted p-values are reported.

### RNA Extraction, cDNA Synthesis and qPCR Assay and Analysis

Total RNA was extracted from I3 neurons using TRIzol reagent (Thermo Fisher Scientific) following the manufacturer’s protocols. RNA was quantified through Cytation 5 Agilent BioTek Cytation 5 Cell Imaging Multimode Reader (A260/A280 ratio ≥ 1.9 required). cDNA was synthesized from 500 ng–1 µg of total RNA using the SuperScript IV (Thermo Fisher Scientific) following the manufacturer’s instructions. Quantitative PCR was performed on a QuantStudio 5 Real-Time PCR System using Power SYBR Green PCR Master Mix (Applied Biosystems) for selected targets. Primer sequences are provided in **Table 2**. Relative expression was calculated by the 2−ΔΔCt method using GAPDH and/or TUJ1 as reference genes. All qPCR assays were performed in technical triplicates.

**Table 2.**
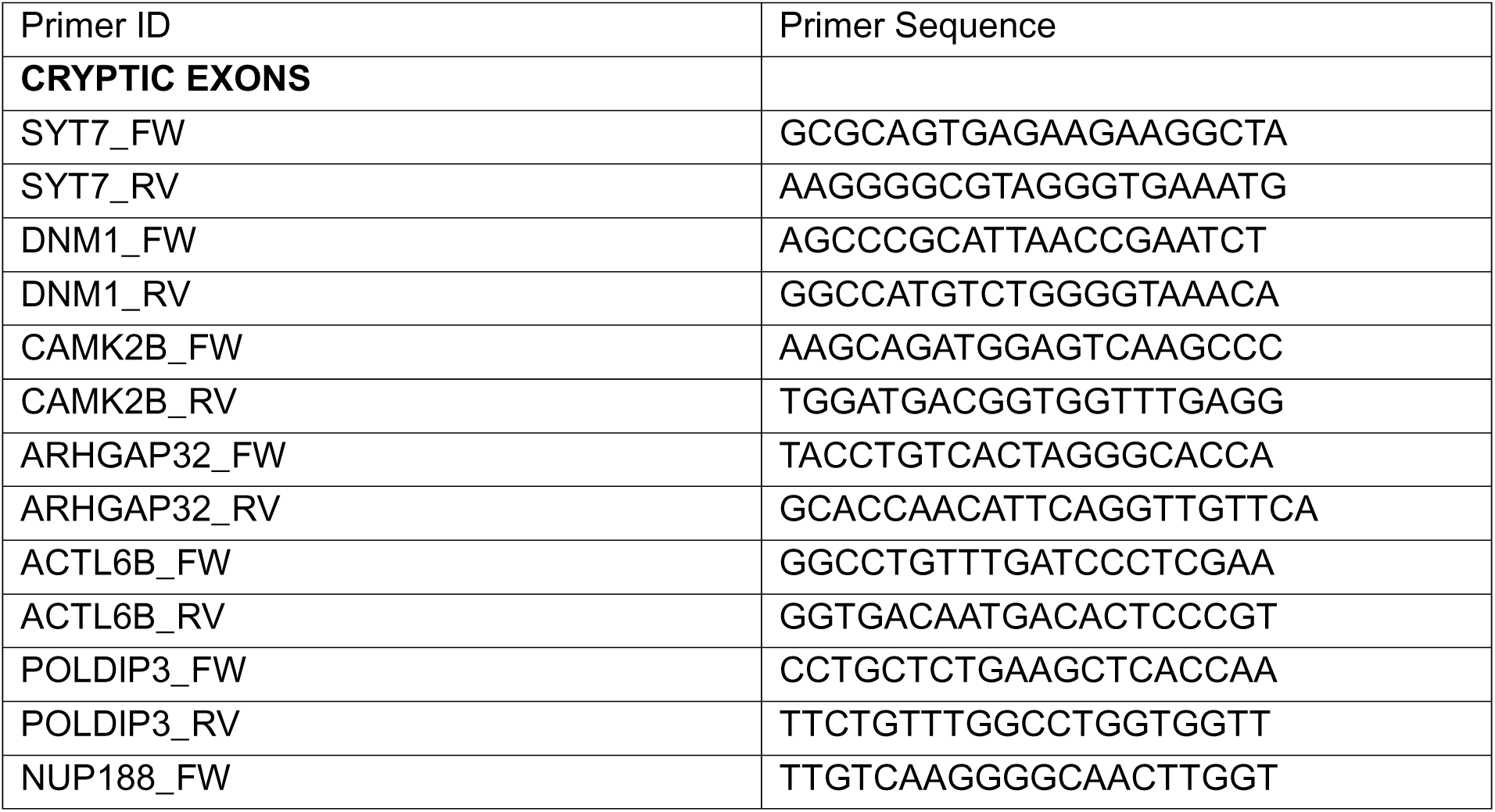
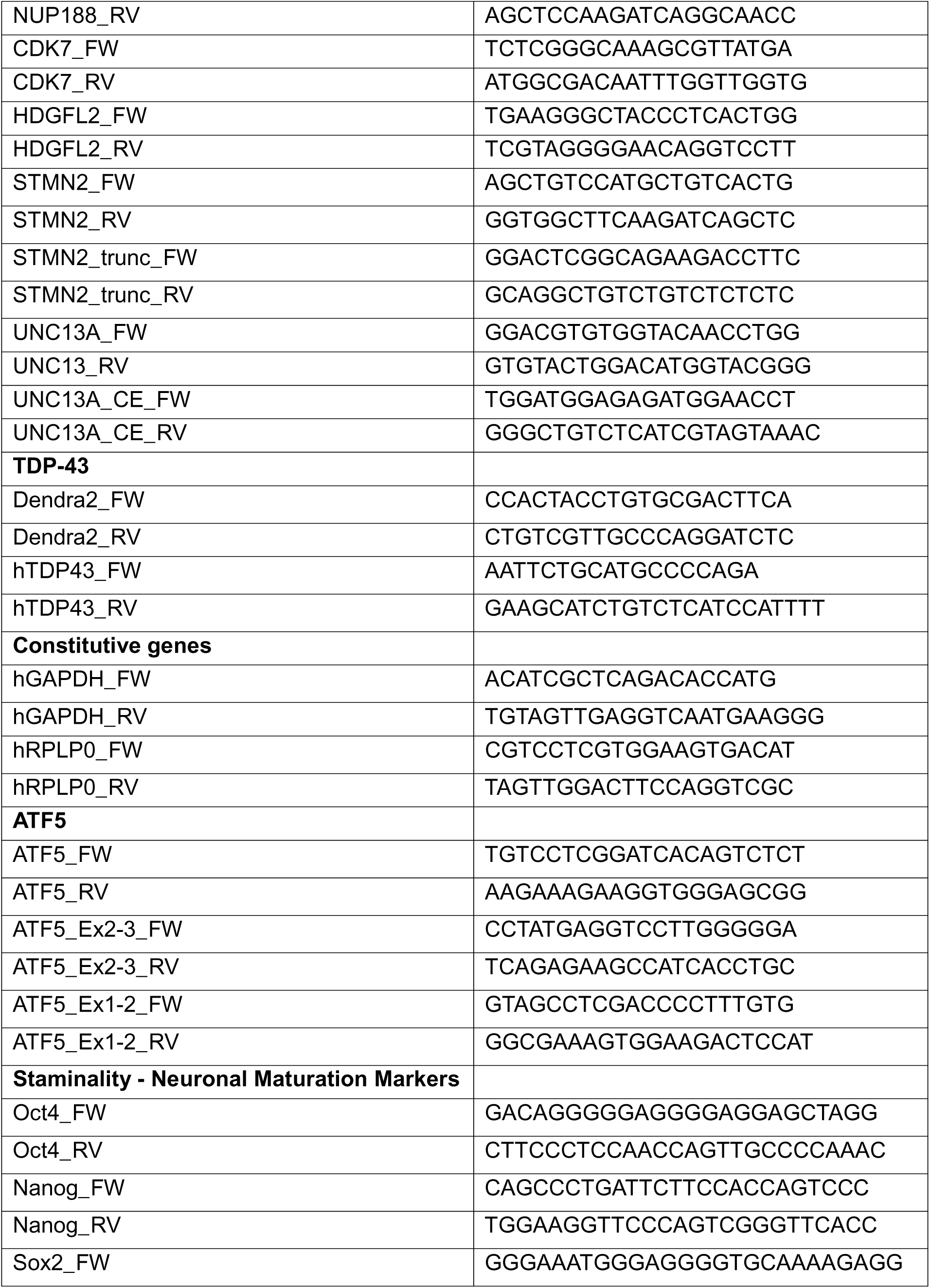
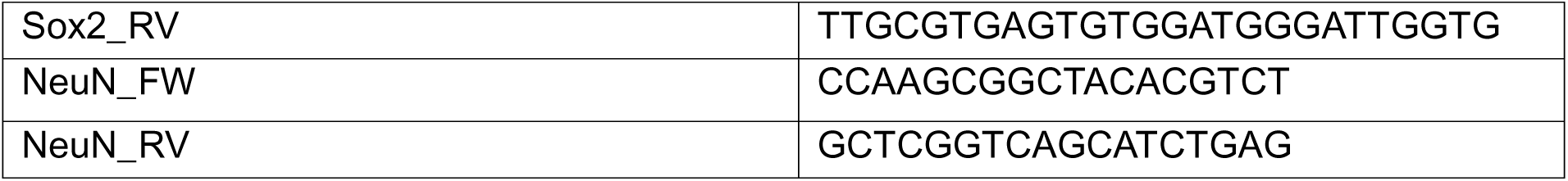
qPCR primers sequence.

### PCR (Skiptic Exon, Cassette)

PCR was conducted using MyTaq Red Mix (Meridian Bioscience) according to the manufacturers protocol. The primer sequences, product sizes of full length and skipped exon products, and reaction specific PCR annealing temperatures are as follows. The skiptic exon primers used for our study were previously generated and described^35^. PCR products were resolved on 2.5% agarose gels with a 100bp DNA ladder (Thermo Scientific) for size visualization. Positive control cDNA from TDP-43 overexpressing HEK293 cells was kindly provided by^35^.

### MEA

Neuronal activity was assessed using multielectrode array (MEA) recordings. Cortical neurons were plated (120,000 cells/well) onto MEA plates coated with PLO and laminin (as previously described). Spontaneous electrical activity was recorded longitudinally under physiological culture conditions every 2-3 days from DIV 10 onward. Key parameters including mean firing rate, burst frequency, burst duration, inter-spike interval, and network synchrony were recorded.

### INCUCYTE

Neuronal morphology was monitored longitudinally using the Incucyte live-cell imaging platform in brightfield mode. Cortical neurons were plated (45,000 cells/well) in a 24 well plate. Automated image acquisition was performed every 8 hours under physiological culture conditions from DIV10 to DIV 40. Quantitative analyses of neurite outgrowth, neurite branching complexity was performed by Neurotrack analysis module.

### OCR and ECAR Monitoring

Oxygen consumption levels were continuously monitored by Resipher platform (Lucid Scientific) on live neurons. Neurons were plated on a flat-bottom 96-well plate (80,000 cells/well) in 100 μL of cortical maturation media. At DIV10, oxygen consumption levels were continuously monitored for 50 hours in a constant-temperature incubator at 37C and 5% CO2. Oxygen consumption rate(OCR) was automatically calculated by the Resipher software based on an oxygen diffusion kinetics model. Since neurons are postmitotic cells, the two lines were plated at the same density, and no normalization on total protein was performed for this experiment. OCR and extracellular acidification rate (ECAR) were simultaneously measured using DolphinQ system (Leadgene Biosolutions). Neurons were plated in cortical maturation media on specific sensor plates (80,000 cells/well) coated with PLO and laminin, as previously described. At DIV10, the DolphinQ mixing lid was placed on the sensor plate, and that assembly was then positioned atop the plate holder tray. 40 mL of ddH₂O were added to the water tray surrounding the plate. O2 concentration and pH were continuously monitored from DIV10 to DIV 17. OCR and ECAR were inferred from O2 concentration and pH by the DolphinQ software, version 2.6.1.1.

### Nucleus, Cytoplasmic and Mitochondrial Fractionation

Mitochondrial, nuclear and cytosolic fractionation was performed according to the protocol described by Bryant et al.^49^. Neurons were washed with PBS and then resuspended in 100 µL of Digitonin Lysis Buffer (50 mM HEPES pH 7.4, 150 mM NaCl, 18 µg/mL digitonin, supplemented with protease inhibitors), incubated on a rotating platform for 10 minutes at 4 °C, and centrifuged at 950 × g for 5 minutes at 4 °C. The supernatant was collected and transferred to a fresh tube as the cytosolic fraction, while the pellet was kept on ice for further processing. To remove debris from the cytosolic fraction, the collected supernatant was centrifuged at 17,000 × g for 5 minutes at 4 °C. The resulting supernatant was saved as the final cytosolic fraction, and the pellet was discarded. The pellet retained from the first centrifugation was washed three times with cold PBS, centrifuged at 950 × g for 3 minutes at 4 °C to remove residual cytosolic proteins. After washing, the pellet was lysed in 100 µL of NP-40 Lysis Buffer (50 mM Tris-HCl pH 7.5, 150 mM NaCl, 1 mM EDTA, 1% NP-40, 10% glycerol, balance H₂O) supplemented with protease and phosphatase inhibitors and incubated on ice for 10 minutes. The lysate was then centrifuged at 21,000 × g for 10 minutes at 4 °C, and the resulting supernatant was collected as the mitochondrial fraction. The pellet was washed twice with cold PBS (centrifugation at 21,000 × g, 3 minutes, 4 °C) and resuspended in 100 µL of SDS Lysis Buffer (10% SDS, 1 M Tris-HCl pH 8, nuclease-free water). Samples were boiled at 95 °C for 15 minutes, sonicated for 5 minutes, and centrifuged at maximum speed for 5 minutes. The supernatant was collected as the nuclear fraction. Samples were prepared for Western Blot analysis by mixing 12.5 µL of each fraction with 2.5 µL of loading dye (included in the total 15 µL per lane) and boiled at 95 °C for 10 minutes. Fraction purity was confirmed by immunoblotting with compartment-specific markers: Lamin A/C (nuclear), GAPDH (cytoplasmic), and COXIV (mitochondrial).

### Western Blot

Cells lysates were separated by SDS-PAGE on 4–12% Mini-PROTEAN® TGX™ Precast Gels (Bio-Rad) and transferred to nitrocellulose membranes (0.45 µm, Millipore) using a wet or semi-dry transfer system (Bio-Rad). Membranes were blocked in 5% non-fat dry milk or 5% BSA in TBST for 1 hour at room temperature and incubated overnight at 4°C with primary antibodies: anti-TDP-43 (1:1000, Proteintech), anti-COXIV (1:2000, Cell Signaling Technology), anti-Lamin A/C (1:1000, Abcam), anti-GAPDH (1:3000, Proteintech), anti-LC3 (1:1000, Signa-Aldrich), and anti-α-Tubulin (1:1000, Proteintech). HRP-conjugated secondary antibodies (anti-rabbit or anti-mouse IgG, 1:5000–1:10000, Thermo Scientific) were used for signal detection by enhanced chemiluminescence (SuperSignal West Femto, Thermo Fisher Scientific). Membranes were imaged on a ChemiDoc MP. Total protein loading was assessed by Ponceau S staining prior to antibody incubation. Band intensities were quantified using ImageLab.

### Soluble–Insoluble Extraction

Soluble and insoluble protein extraction was performed following a protocol adapted from Wagner et al^50^. Cells were detached from multi-well plates, pelleted, washed to remove medium/PBS, and stored at −80 °C until processing. Cell pellets were resuspended in 300 µL of RIPA buffer supplemented with protease and phosphatase inhibitors and lysed by sonication. Lysates were incubated on ice for 15 min, and 30 µL of each sample were mixed with Laemmli buffer containing 5% β-mercaptoethanol and boiled at 95 °C for 10 min to obtain the Input fraction. The remaining lysate (150 µL) was centrifuged at 20,800 g for 20 min at 4 °C. 130 µL of the supernatant were collected, mixed with Laemmli buffer, and boiled to generate the soluble fraction. The pellet was washed with 1 mL of RIPA buffer (20,800 g, 4 °C, 15 min), and the supernatant was discarded. Pelleted proteins were then resuspended by sonication (10 s, 25% amplitude) in 56.25 µL of resolubilization buffer (7 M urea, 2 M thiourea, 4% CHAPS, 30 mM Tris, pH 8.5). Insoluble debris was removed by centrifugation (20,800 g, RT, 15 min), and 40 µL of the resulting supernatant were mixed with Laemmli buffer and boiled to obtain the insoluble fraction.

### Immunofluorescence and Confocal Microscopy

Neurons were plated on poly-L-ornithine/laminin-coated glass 24 well-plate. At the designated time points, cells were fixed with 4% paraformaldehyde (PFA) in PBS for 15 minutes at room temperature, permeabilized and blocked with 0.1% Triton X-100, 1% FBS and 0.1% BSA in PBS for 1 hour at room temperature. Primary antibodies were diluted in 0.01% BSA in PBS and incubated overnight at 4°C: anti-TDP-43 (1:500), anti-MAP2 (1:2000, Novus Biological), anti-G3BP1 (1:1000, Proteintech), anti-NANOG (1:200 R&D System), anti-OCT4 (1:250, Santa Cruz), anti-SOX2 (1:200 EMD Millipore). Secondary antibodies conjugated to Alexa Fluor 488, 546, or 647 (Thermo Fisher Scientific, 1:1000) were incubated for 1 hour at room temperature protected from light. Nuclei were counterstained with Hoechst 33342 (1 µg/mL). Samples were then kept in PBS. Images were acquired on a Nikon A1 confocal microscope using 20× (dry) or 63× (oil immersion, NA 1.4) objectives. All acquisition settings (laser power, gain, pinhole) were kept constant across conditions within each experiment.

### Image Analysis (Nuclear TDP-43, TDP-43 Aggregates, SGs Analysis)

Image analysis was performed using the Nikon NIS-Elements software. For quantification of nuclear TDP-43 intensity, nuclei were segmented based on the BFP signal, and the mean fluorescence intensity of the Dendra2 or total TDP-43 channel within the nuclear mask was measured.

For aggregate quantification, nuclear and cytoplasmic aggregates were identified by thresholding the Dendra2 signal. Aggregate number was quantified either within the mask drawn around each nucleus or within the cytoplasmic mask generated by subtracting the nuclear mask from an enlarged nuclear mask.

For stress granule analysis, G3BP1-positive puncta were segmented by thresholding the G3BP1 channel, and the mean fluorescence intensity of Dendra2 within G3BP1 masks was measured and reported as Dendra2 SG MFI. The number of stress granules was quantified in the cytoplasm using the approach described above.

For neurite length quantification, neurons were traced using Nikon NIS-Elements, and the total neurite length per field was recorded. A minimum of 50–100 cells per condition were analyzed in each experiment, and all experiments were independently repeated at least three times.

### Statistical Analysis

Statistical analyses were performed in GraphPad Prism (version 9 or later) or R (version 4.x). Data are presented as mean ± SEM unless otherwise indicated. For comparisons between two groups, unpaired two-tailed Student’s t-tests were used. For comparisons across multiple groups or conditions, One-Way or Two-Way ANOVA followed by Tukey’s post-hoc test was applied as appropriate.

## Figure Legends

**Supplementary Figure 1. Generation and characterization of isogenic TDP-43 A315T Dendra2-tagged iPSC lines.**

(a) Schematic overview of the CRISPR-Cas9 gene editing strategy used to introduce the A315T point mutation and an in-frame Dendra2 fluorescent tag into the C-terminus of the endogenous TDP-43 locus, generating two isogenic iPSC lines: TDP-43 A315T_Dendra2 and TDP-43 WT_Dendra2.

(b) Sanger sequencing chromatograms of TDP-43 exon 6 confirming the GCG→ACG (Alanine→Threonine) nucleotide substitution in the TDP-43 A315T line relative to the TDP-43 WT sequence.

(c) Karyotype analysis of TDP-43 WT (left, green box) and TDP-43 A315T (right, red box) iPSC lines confirming normal diploid chromosomal content (46, XY) with no detectable chromosomal abnormalities.

(d) Representative brightfield images of iPSC colonies stained for Alkaline Phosphatase activity, a marker of pluripotency, in TDP-43 WT (left) and TDP-43 A315T (right) human induced pluripotent lines. Scale Bar = 100µm.

(e) Representative immunofluorescence images showing nuclear NANOG expression (magenta) in TDP-43 WT and TDP-43 A315T iPSCs colonies. Scale Bar = 25µm.

(f) Representative immunofluorescence images of TDP-43 WT (top row) and TDP-43 A315T (bottom row) iPSCs co-stained for DAPI (blue), OCT4 (red), SOX2 (magenta), and Dendra2 (green), with merged channel overlay. Scale Bar = 10µm.

**Supplementary Figure 2. Derivation and characterization of I3 cortical neurons from isogenic iPSC lines.**

(a) Schematic of hNGN2 cassette insertion into iPSCs lines through piggyback system and subsequent positive clones selection through antibiotic selection and flow cytometry based sorting.

(b) Representative brightfield images of TDP-43 WT and TDP-43 A315T iPSC-derived I3 neurons after puromycin selection (+plasmids) versus a no-plasmid puromycin control. Scale Bar = 100μm.

(c) Representative FACS gating strategy for sorting BFP- and Dendra2-positive iPSC-derived neurons in TDP-43 WT (top, green box) and TDP-43 A315T (bottom, red box) lines.

(d) PCR validation of NGN2-PiggyBac insertion. Agarose gel showing the expected 320 bp band from TRE (3X) -NGN2 locus in both TDP-43 WT and TDP-43 A315T lines.

(e) Representative brightfield images of iPSC colonies from TDP-43 WT and TDP-43 A315T lines stained for Alkaline Phosphatase activity. Scale Bar = 100µm.

(f) Representative immunofluorescence images of iPSC colonies from TDP-43 WT and TDP-43 A315T lines co-stained for SOX2 (red/magenta), OCT4 (red), Dendra2 (green), and BFP (blue). Scale Bar = 100µm.

(g–j) qRT-PCR analysis of SOX2 (g), OCT4 (h), NANOG (i) and NEUN (j) marker expression in TDP-43 WT and TDP-43 A315T iPSCs clones pre (iPS) and post (iPS_i3) cassette insertion. Data presented as mean ± SEM, n = 3 independent replicates.

(k) Schematic diagram summarizing the timeline of I3 neuron maturation. BFP/Dendra2-double positive cells progress through neural induction (Day 1), immature neuron stages (Days 1–5), and mature into cortical neurons (Day 21) following doxycycline-induced NGN2 expression.

**Supplementary Figure 3. Raw Ct values for qRT-PCR assays.**

(a) RT-qPCR analysis (Raw Ct values) for UNC13A, STMN2, GAPDH, TUJ1, STMN2-TRUNC, and UNC13A CE in TDP-43 WT (green) and TDP-43 A315T (red) neurons treated with the NMD inhibitor NMDi14 (5 µM for 24 hours or 50 µM) for 48 hours) or DMSO vehicle. Data are presented as mean ± SEM. n = 3-6 independent replicates. *p<0.05, Student T-test

Supplementary Figure 4. LC3 lipidation is increased in TDP-43 A315T neurons, indicating elevated autophagic flux.

(a) Representative immunoblot of total cell lysates from TDP-43 WT and TDP-43 A315T I3 neurons probed with anti-LC3 antibody (αLC3) and anti-alpha-Tubulin (αTUB) as a loading control.

(b–d) Quantification of LC3 isoforms normalized to Tubulin. (b) LC3-I/Tubulin ratio; (c) LC3-II/Tubulin ratio; (d) LC3-II/LC3-I ratio Data presented as mean ± SEM, n = 3 independent experiments. *p<0.05, **p<0.01 Student T-test.

