## Supplementary figures and images for "A Human Neuronal Cell Model of Endogenous TDP-43 A315T Reveals Altered Protein Dynamics and Disease-Relevant Cellular Dysfunction"

### Suppl_Fig

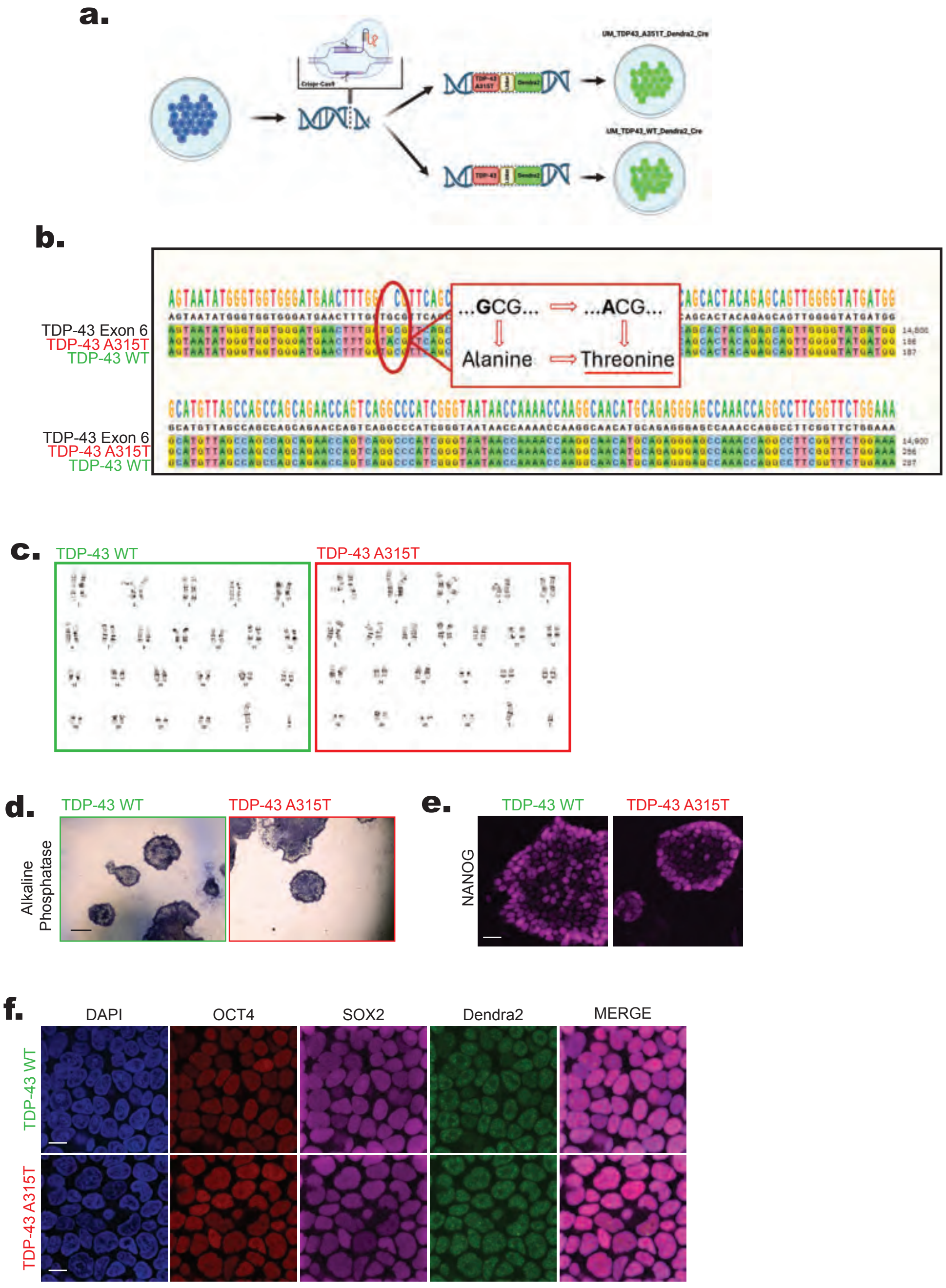


Supplementary Figure 1

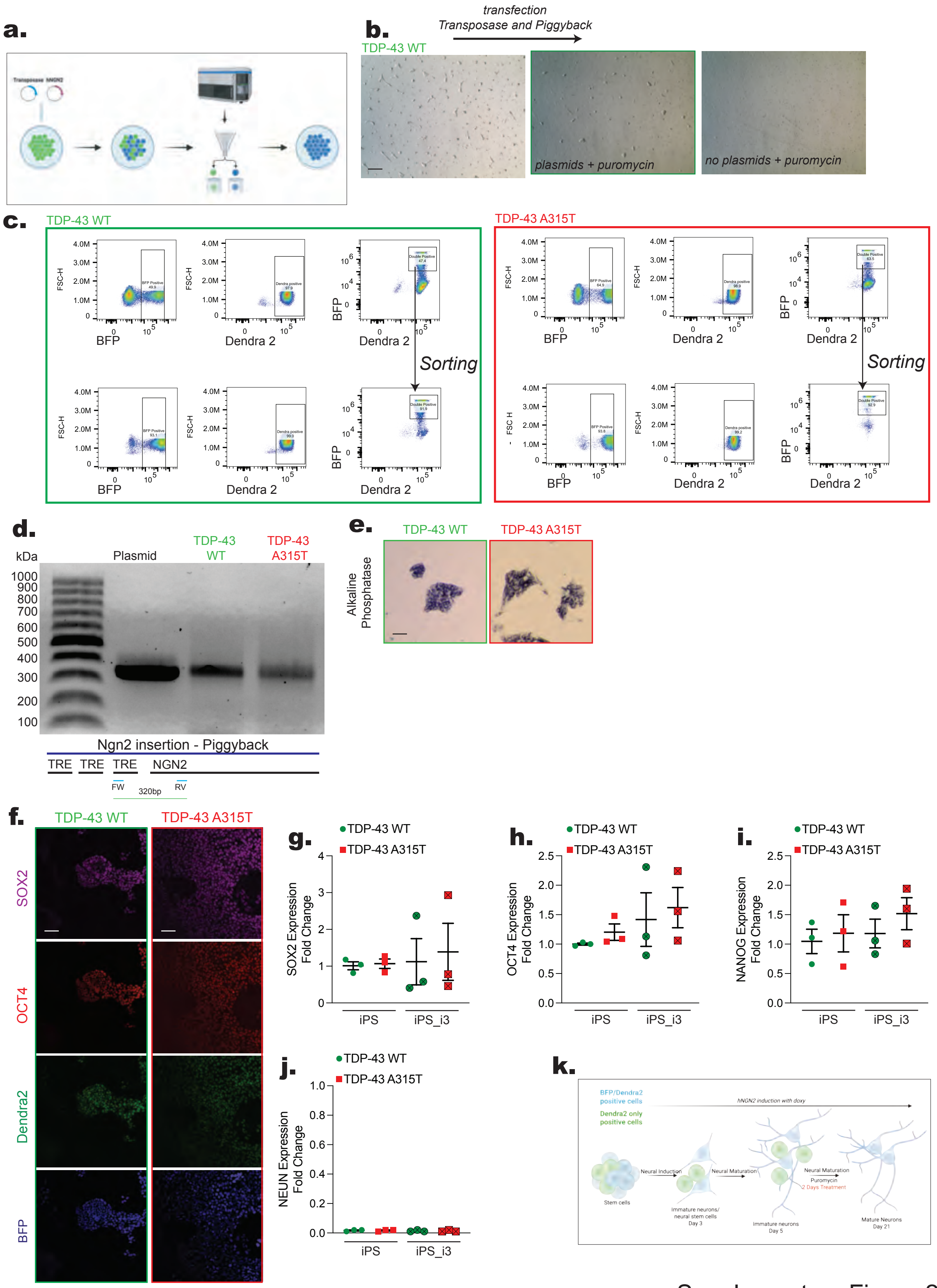


Supplementary Figure 2

**a.**

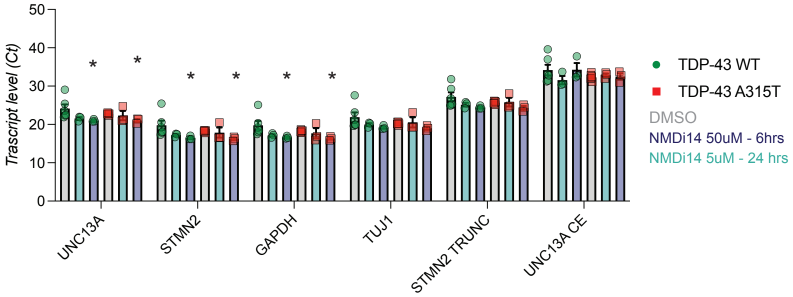


Supplementary Figure 3

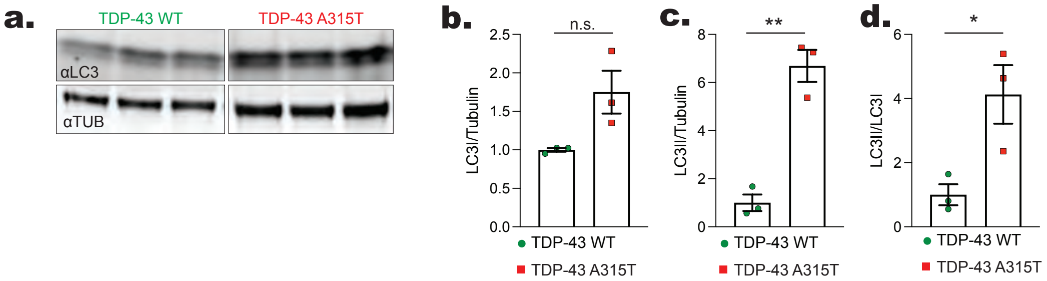


Supplementary Figure 4
